# Solving the spike sorting problem with Kilosort

**DOI:** 10.1101/2023.01.07.523036

**Authors:** Marius Pachitariu, Shashwat Sridhar, Carsen Stringer

## Abstract

Spike sorting is the computational process of extracting the firing times of single neurons from recordings of local electrical fields. This is an important but hard problem in neuroscience, complicated by the non-stationarity of the recordings and the dense overlap in electrical fields between nearby neurons. To solve the spike sorting problem, we have continuously developed over the past eight years a framework known as Kilosort. This paper describes the various algorithmic steps introduced in different versions of Kilosort. We also report the development of Kilosort4, a new version with substantially improved performance due to new clustering algorithms inspired by graph-based approaches. To test the performance of Kilosort, we developed a realistic simulation framework which uses densely sampled electrical fields from real experiments to generate non-stationary spike waveforms and realistic noise. We find that nearly all versions of Kilosort outperform other algorithms on a variety of simulated conditions, and Kilosort4 performs best in all cases, correctly identifying even neurons with low amplitudes and small spatial extents in high drift conditions.

## Introduction

Classical spike sorting frameworks require a sequence of operations, which can be categorized into preprocessing, spike detection, clustering and postprocessing. Modern approaches have improved on these steps by introducing new algorithms. Some frameworks [1–3] took advantage of new clustering algorithms such as density-based approaches [4] or agglomerative approaches using bimodality criteria [5]. In contrast, the original Kilosort [6] used a simple clustering approach (scaled K-means), but combined two steps of the pipeline into one (spike detection + clustering = template learning) and added an extra matching pursuit step for detecting overlapping spikes, sometimes referred to as solving the “collision problem” [7].

An important consideration for these early modern algorithms was the requirement for additional human curation, as the clustering results were imperfect in many applications. Thus, algorithms like Kilosort biased the clustering process towards “over-splitting”, producing more clusters than the number of real units in the data, so that human curation would consist primarily of merges, which are substantially easier to perform than splits. To facilitate human curation of the automated results, a modern graphical user interface called Phy was developed, which is now used for visualization by several of the most popular frameworks including all versions of Kilosort [8].

Why was human curation still necessary for these early modern methods? One of the main reasons was the non-stationary nature of data from real experiments. The electrical field of a unit sampled by a probe, called a spike waveform, should be fixed and reproducible across long time periods. Yet in many experiments, the shape of the waveform appeared to change over the course of hours, and sometimes much faster. The main reason for these changes was identified as vertical probe movement or “ drift”, using high-density electrodes [9]. Drift is primarily caused by factors such as tissue relaxation after probe insertion and animal movements during behavior. Correcting for this drift resulted in substantial improvements in spike sorting performance. Kilosort2 used a “drift tracking” approach for this, while Kilosort2.5 developed a standalone drift correction method that directly modified the voltage data to shift certain channels up or down by appropriate distances (see Methods for drift tracking, and Methods in [9] for drift correction). The drift correction step has been inherited by all Kilosort versions since 2.5.

The main goal of this paper is to describe the development of Kilosort4 and demonstrate its performance. Some of the algorithmic steps in Kilosort4 are inherited from previous versions (i.e. drift correction), while others build on top of previous versions (i.e. template deconvolution), while others are completely new (i.e. the graph-based clustering approach). Except for drift correction, which was previously described in detail [9], the other algorithmic steps are not described in the literature, and we add detailed descriptions in the Methods (see Table 1 for an overview). We also developed a new simulation-based framework for benchmarking spike sorting algorithms, which uses several realistic drift patterns and dense electrical fields inferred from real experiments. We show using the benchmarks that Kilosort4 performs very well and outperforms all other algorithms across a range of conditions.

**Table 1:** The evolution of kilosort. New features added after Kilosort 1 are bolded in the version they were first introduced. ^1^ described in Pachitariu et al, 2016 *bioRxiv*, ^2^ described in Steinmetz et al, 2021 *Science*, ^∗^ described in this paper.

| algorithms | language | pre-processing |  | template deconvolution |  |  |  | clustering & post-processing |  |  |
| --- | --- | --- | --- | --- | --- | --- | --- | --- | --- | --- |
|  |  | filtering & whitening | drift correction | template deconvolution | template learning | deconv. during learning | new templates from residual | clustering | splits | merges |
| Kilosort 1 (2016) | matlab +CUDA | yes<sup>1</sup> | — | yes<sup>1</sup> | scaled k-means<sup>1</sup> | — | — | — | — | — |
| Kilosort 2 (2018) | matlab +CUDA | yes | — (only **tracking\***) | yes | scaled k-means | **yes\*** | **threshold crossing\*** | — | **bimodality pursuit\*** | **yes\*** |
| Kilosort 2.5 (2020) | matlab /python +CUDA | yes | **yes**<sup>2</sup> | yes | scaled k-means | yes | threshold crossing | — | bimodality pursuit | yes |
| Kilosort 3 (2021) | matlab +CUDA | yes | yes | yes | **recursive pursuit\*** | — | — | **recursive pursuit\*** | yes | yes |
| Kilosort 4 (2023) | python +pytorch | yes | yes | yes | **graph clustering\*** | — | — | **graph clustering\*** | **merging tree\*** | yes |

## Results

To be able to process the large amounts of data from modern electrophysiology, all versions of Kilosort are implemented on the GPU. Kilosort4 is the first version fully implemented in python and using the pytorch package for all its functionality, thus making the old CUDA functions obsolete [10, 11]. Pytorch allows the user to switch to a CPU backend which may be sufficiently fast for testing on small amounts of data but is not recommended for large-scale data. All versions of Kilosort take as input a binary data file, and output a set of “.npy” files that can be used for visualization in Phy [8]. To set up a Kilosort4 run, we built a pyqtgraph GUI which replicates the functionality of the Matlab GUI, and can assist users in debugging due to several diagnostic plots and summary statistics that are displayed [12] (Figure S1).

The preprocessing step in all versions of Kilosort includes temporal filtering and channel whitening (see Methods). These linear operations reduce the strong spatiotemporal correlations of the electrical background in the brain, which is mainly formed by the electrical discharge of units that are too far from the probe to be identified as single units. This step is accelerated in Kilosort4 through the use of explicit convolutions in place of a Butterworth filter. Drift correction is an additional preprocessing step that was introduced in Kilosort2.5 and maintained in all subsequent versions (see Methods of [9]). Unlike previous versions, Kilosort4 no longer needs to generate an intermediate file of processed data, as all preprocessing operations are fast enough to be performed on-demand.

### Template deconvolution

We refer to the spike detection and feature extraction steps jointly as “template deconvolution”. This module requires a set of templates which correspond to the average spatiotemporal waveforms of neurons in the recording. The templates are used in the matching pursuit step for detecting overlapping spikes [6]. A template deconvolution step has been used in all versions of Kilosort, but the details of the template learning have changed (see Methods). In Kilosort 3 and 4, the template deconvolution serves an extra role as a feature extraction method with background correction.

The template deconvolution pipeline has the same format for both Kilosort 3 and 4 (Figure 1a). A set of initial spike waveforms are extracted from preprocessed data using a set of universal templates (Figure 1b,c). The features of these spikes are then clustered, using either the recursive pursuit algorithm from Kilosort3 (see Methods), or the graph-based algorithm from Kilosort4 (described in Figure 2). The centroids of the clusters are the “learned templates”, which are then aligned temporally (Figure 1d). The templates are compared to each other by cross-correlation and similar templates are merged together to remove duplicates. The learned templates are then used in the matching pursuit step, which iteratively finds the best matching templates to the preprocessed data and subtracts off their contribution. The subtraction is a critical part of all matching pursuit algorithms and allows the algorithm to detect spikes that were overlapped by the subtracted ones. The final reconstruction of the data with the templates is shown in Figure 1e. The residual is the difference between data and reconstruction, and can be informative if the algorithm fails to find some units (Figure 1f).

**Figure 1:**
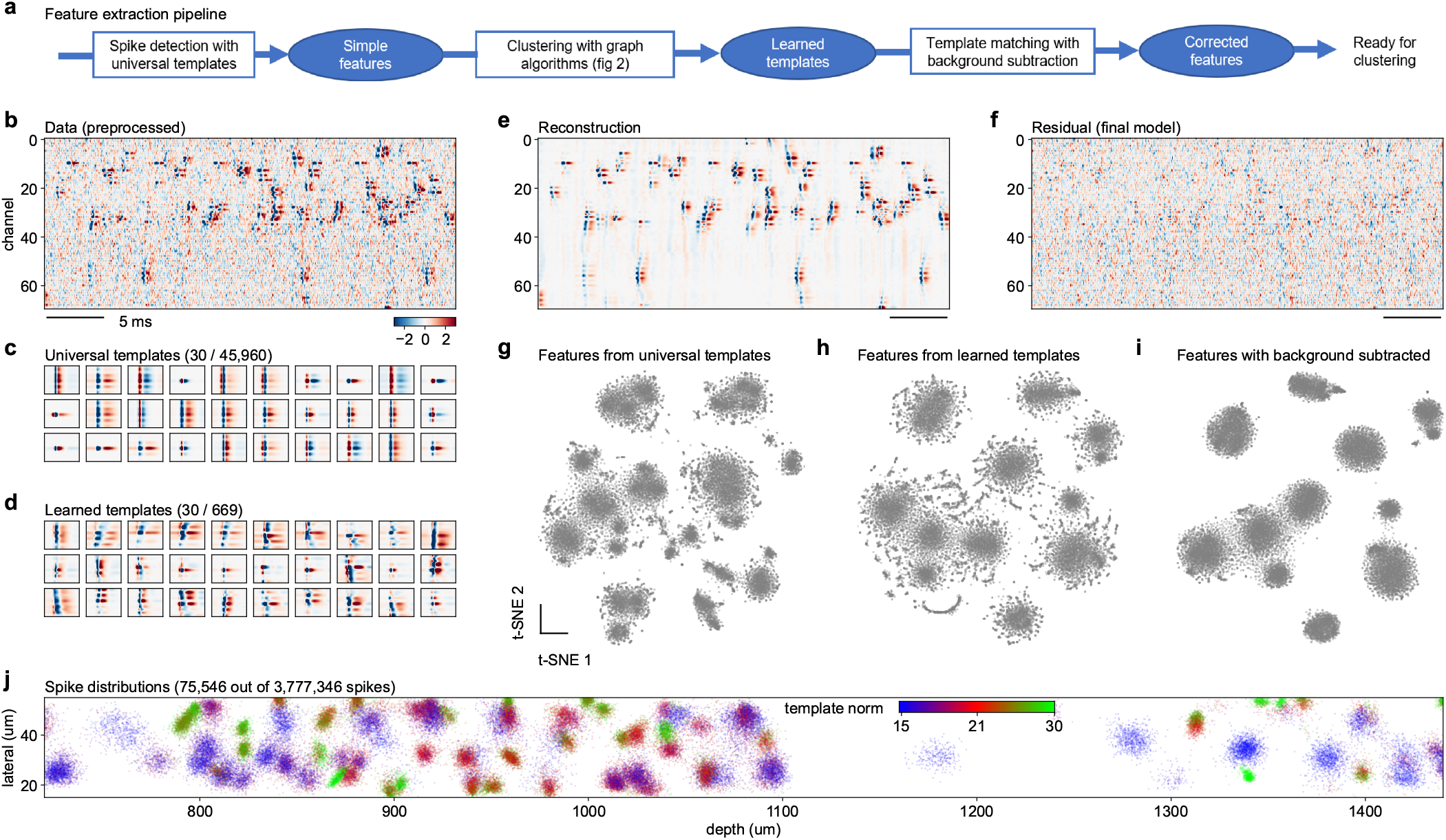
Spike detection and feature extraction. **a**, Schematic of the pipeline for detecting spikes and extracting spikes features. **b**, Short segment of preprocessed data over 70 channels and 1,000 timepoints (data from [13]). **c**, Example universal templates centered at a single position on the probe. Templates are repeated at 1536 positions for a Neuropixels probe. **d**, Example learned templates centered at different positions on the probe. **e**, Reconstruction of the data in **b** based on the inferred templates and spike times. **f**, Residual after subtracting the reconstruction from the data. **g-i**, t-SNE visualization of spike features from a 40µm segment of the probe. Spike features were extracted using either universal templates (**g**) or learned templates without (**h**) or with (**i**) background subtraction. **j**, Spatial distribution of a subset of the final extracted spikes colored by their template norm.

**Figure 2:**
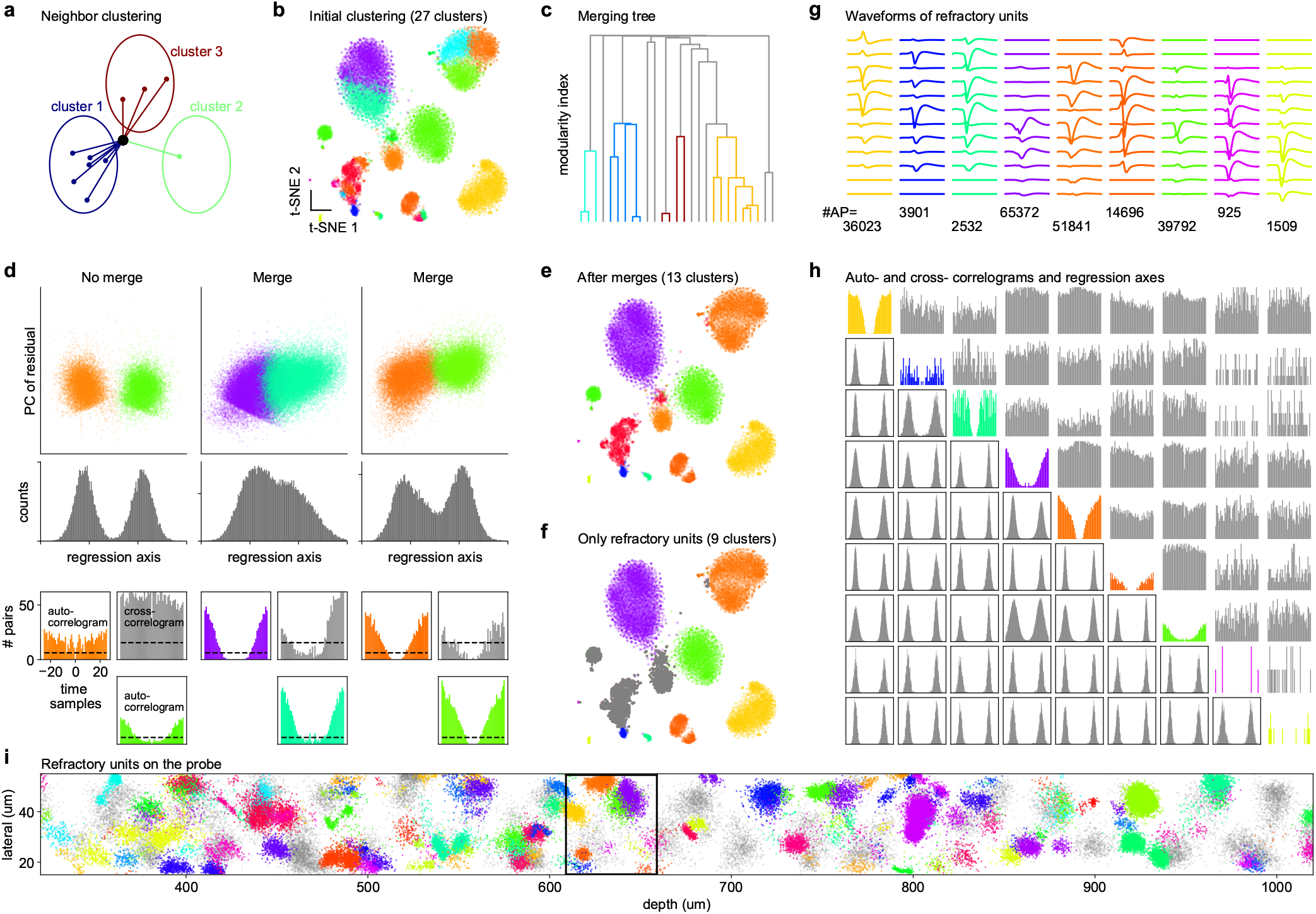
Clustering spike with graph-based methods. **a**, Illustration of the iterative re-assignment process. At each iteration, a node is assigned to the cluster which has most of its neighbors. A penalty is used to compensate for larger clusters having more neighbors. This process is initialized with 200 clusters obtained from K-means++ [14] and converges to a smaller set of clusters in tens of iterations. **b**, Example clustering produced by this process overlaid on a t-SNE visualization (data from [15]). **c**, Merging tree formed by merging clusters according to the modularity cost function. Colored branches correspond to merges that were accepted. The tree is traversed from top to bottom to make split/merge decisions. **d**, Criteria for performing a merge/split decision in the merging tree: (top) projection across the regression axis has to be bimodal, (bottom) cross-correlogram of spike times cannot be refractory (dashed line indicates approximate refractory criterion). **e**, Final result of the clustering algorithm after merges. **f**, Same as **e** with non-refractory units grayed out. **g**, Average waveforms of units with refractory periods and the total number of spikes in each cluster. **h**, (diagonal) Auto-correlograms; (below diagonal) projection on regression axes; (above diagonal) cross-correlograms. **i**, Subset of spikes colored by their final assigned clusters. Non-refractory units shown in gray.

This template learning step from Kilosort 3/4 is different from the one in Kilosort1 (see [6]), and both are different from the equivalent step in Kilosort 2/2.5 (see Methods). Furthermore, the templates of Kilosort1 are in one-to-one correspondence with the final inferred units which are exported for manual curation. This correspondence is weaker in Kilosort 2/2.5, because a post-processing step is used to perform splits and merges on these templates (see Methods). Finally, in Kilosort 3/4 these templates are completely discarded after being used to extract spikes. This is because more powerful clustering algorithms can be applied to the spike features once they have been extracted with template deconvolution. The “corrected” or deconvolved features have three additional properties compared to the features detected with universal templates or more generally detected with any classical threshold crossing method: 1) they contain all, or a majority of spikes from the clustered units, even the ones that are overlapped by larger, bigger spikes; they group spikes together by templates, which can be used to more precisely assign spikes to their best channels for batched clustering across channels; 3) they can be computed after subtraction of the background produced by all other spikes (Figure 1e).

These properties have a substantial effect on the features, allowing for better clustering. Figure 1g-i show the t-SNE embeddings of three different sets of features from spikes detected over a 40um stretch of a Neuropixels probe. The features computed with the learned templates with background subtraction (Figure 1i) are embedded as more uniform, Gaussian-like clusters. Without background subtraction, each cluster is surrounded by a patterned envelope of points due to the contribution of overlapping spikes, and these patterns can be easily mistaken for other clusters (Figure 1g,h). The visualization in Figure 1i can be used to get an impression of a small section of the data without performing any clustering at all. To visualize the distribution of spikes over a larger portion of a probe, we plot a subset of spikes at their inferred XY positions (Figure 1j). The spikes are colored according to amplitudes, which tends to be uniform for spikes from the same unit.

### Graph-based clustering with merging trees

We developed two new clustering algorithms for spike features extracted by template deconvolution. In Kilosort3, we developed an algorithm that uses a recursive application of the bimodality pursuit algorithm from Kilosort2, which in turn had been developed to automatically find potential splits within clusters (see Methods). In Kilosort4 we developed a graph-based clustering method. This approach first constructs a graph of points connected to their nearest neighbors in Euclidean space, then constructs a cost function from the graph properties to encourage the clustering of nodes. A popular cost function is “modularity”, which counts the number of graph edges inside a cluster and compares them to the expected number of edges from a disorganized, unclustered null model [16]. Well-known implementations of modularity optimization are the Leiden and Louvain algorithms [17, 18]. Applied directly to spike features, these established algorithms fail in a few different ways: 1) difficulty partitioning clusters with very different number of points; 2) very slow processing speed for hundreds of thousands of points; 3) cannot use domain knowledge to make merge/split decisions.

To remedy these problems, we developed a new graph-based algorithm, in which the clusters are defined as the stationary points of an iterative neighbor reassignment algorithm based on the modularity cost function (Figure 2a and see Methods). This method allowed us to find more of the small clusters compared to a straightforward application of the Leiden algorithm (Figure 2b). To improve the processing speed, which typically grows quadratically in the number of data points, we developed a landmark-based version of the algorithm which uses nearest neighbors within a subset of all data points. The application of this algorithm resulted in oversplit clusters, which required additional merges using domain knowledge. To find the best merges efficiently, we used the modularity cost function to construct a “merging tree” (Figure 2c). Potential splits in this tree were tested using two criteria: 1) a bimodal distribution of spike projections along the regression axis between the two sub-clusters (Figure 2d, top), and 2) whether the cross-correlogram was refractory or not (Figure 2d, bottom).

This clustering algorithm was applied to groups of spikes originating from the same 40 µm vertical section of the probe. After all sections were clustered, an additional merging step was performed which tested the refractoriness of the cross-correlogram for all pairs of templates with a correlation above 0.5, similar to the global merging step from previous versions (2/2.5/3). The final results are shown in (Figure 2e). Units that did not have a refractory period are shown grayed out in (Figure 2f); they likely correspond to neurons that were not well isolated. A quick overview of the units identified on this section of the probe shows that all units had refractory auto-correlograms, all pairs of clusters had bimodal projections on their respective regression axes, and all pairs of clusters had flat, non-refractory cross-correlograms (Figure 2h). These properties together indicate that these nine units correspond to nine distinct, well-isolated neurons. These clusters can also be visualized on the probe, in their local contexts (Figure 2i).

### Electrical simulations with realistic drift

To test the performance of Kilosort4 and previous versions, we next developed a set of realistic simulations with different drift patterns. Constructing such a simulation requires knowledge of the dense electric fields of a neuron, because different drift levels sample the electric field at different positions. We obtained this knowledge by sampling neurons from recordings with large drift (Figure 3a) from a public repository of more than 500 Neuropixels recordings from the IBL consortium (Figure 3b). In this repository, we found 11 recordings with large, continuous drift that spanned over at least 40 µm, which is the spatial repetition period of a Neuropixels probe. We separately built two pools of units: one from neurons that were well-isolated and had refractory periods, and one from multi-unit activity which had refractory period contaminations. Drift levels were discretized in 2 µm intervals, and only units with enough spikes in each drift interval were considered. The average waveforms at five positions is shown for a few examples (Figure 3c and Figure S2a,b). To simulate drift, we generated a single average drift trace and additional deviations for each channel to account for heterogeneous drift. Spike trains were generated using shuffled inter-spike intervals from real units. For each simulation, a set of 600 ground-truth neurons were generated in this fashion, with amplitudes drawn from a truncated exponential distribution which matched the amplitudes in real datasets. Another 600 “multi-units” were added with lower amplitudes (Figure 3d). Additional independent noise was added on each channel. The resulting simulation was “un-whitened” across channels using a rotation matrix from real experiments (Figure S2c).

**Figure 3:**
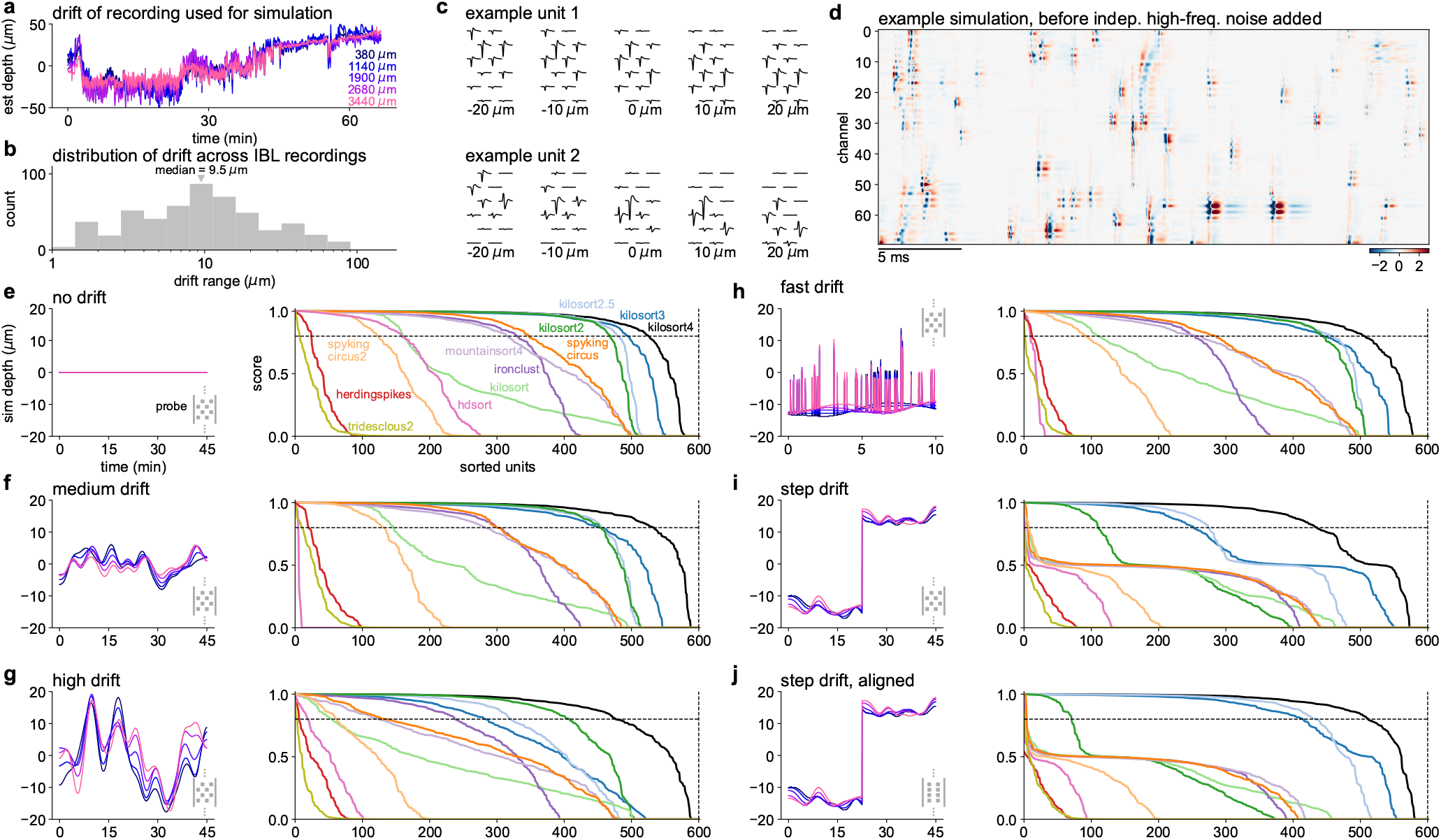
Spike sorting simulations and benchmarks. **a**, Example drift traces at different depths for a recording with large drift from the IBL dataset. **b**, Distribution of drift ranges across all IBL recordings. Drift range was defined as the difference between the 5^th^ and 95^th^ percentile of the median drift across channels. **c**, Waveforms of example units at multiple drift positions. **d**, Segment of a simulated recording with drift over 70 channels and 1,000 timepoints. **e-j**, Accuracy of spike-sorting algorithms on simulations with various drift profiles. Left: simulated drift traces. Right: sorted accuracies for 600 ground truth units from each simulation matched to the results of each algorithm. The accuracy score is defined as 1 - FP - FN, where FP is the false positive rate and FN is the false negative rate (see Methods). **e**, No drift. **f**, Medium drift. **g**, High drift. **h**, Fast drift (10 minutes out of 45 minutes plotted for visibility). **i**, Step drift. **j**, Step drift for a probe with aligned sites.

This simulation framework allowed us to test many algorithms across many simulated experimental conditions [2, 3, 6, 19–23]. All algorithms other than Kilosort4 were run through their respective SpikeInterface wrappers to ensure consistent processing, and parameter adjustments were made in some cases to improve results (see Methods) [24]. The latest algorithm versions as of December 2022 were used in all cases, which are often substantially different from the initial published versions [2, 3]. Results for all conditions are shown in (Figure 3e-j) and quantified in Table 2. All the algorithms had reasonable run times (within 2x the duration of the simulations). The drift conditions we chose were based on patterns of drift identified in the IBL dataset (Figure S3): no drift, medium drift, high drift, fast drift and step drift. The medium drift condition was matched to the median recording from the IBL dataset. The high drift condition had a drift range spanning the entire 40 µm spatial period of the probe, thus sampling all potential shapes of each waveform. The fast drift condition uses drift on the timescale of seconds and sub-seconds, to simulate fast head movements such as during a behavioral task. The step drift condition simulates abrupt changes during an experiment, which are common in the IBL dataset and likely caused by excessive animal movements. This condition also simulates chronic recordings made on different days, where the probe is stationary on each day, but moves in-between days. Since this condition was the most difficult for all algorithms, we also tested whether an aligned sites probe configuration (such as in Neuropixels 2) improves the results.

**Table 2:** Number of correctly identified units in simulations. Number of detected units that matched ground truth units with a 1 - FP - FN score greater than 0.8, for each simulation. Runtime averaged over the 6 simulations of 45 minutes each ± s.e.m. (RTX 3090 + 2x Intel Xeon Gold 6348 + SSD).

| algorithms | no drift | medium drift | high drift | fast drift | step drift | step drift aligned | runtime (minutes) |
| --- | --- | --- | --- | --- | --- | --- | --- |
| Kilosort4 | **526** | **534** | **477** | **506** | **436** | **512** | 25.4 ± 0.7 |
| Kilosort3 | 495 | 454 | 283 | 446 | 253 | 415 | 69.0 ± 2.1 |
| Kilosort2.5 | 479 | 456 | 321 | 460 | 274 | 433 | 24.0 ± 0.5 |
| Kilosort2 | 469 | 454 | 408 | 444 | 110 | 71 | 23.6 ± 0.7 |
| Kilosort [6] | 163 | 147 | 54 | 111 | 7 | 4 | 51.8 ± 0.5 |
| IronClust [1, 19] | 331 | 301 | 239 | 254 | 1 | 1 | 34.9 ± 0.6 |
| MountainSort4 [3] | 316 | 283 | 124 | 262 | 2 | 1 | 82.8 ± 4.4 |
| SpyKING CIRCUS [2] | 348 | 301 | 135 | 285 | 6 | 4 | 65.6 ± 2.1 |
| SpyKING CIRCUS 2 [20] | 123 | 131 | 62 | 93 | 7 | 1 | 32.9 ± 2.3 |
| HDSort [21] | 159 | 5 | 19 | 10 | 1 | 1 | 42.5 ± 1.0 |
| Herding Spikes [22] | 25 | 21 | 6 | 13 | 0 | 0 | 24.7 ± 0.2 |
| Tridesclous2 [23] | 7 | 12 | 3 | 2 | 0 | 1 | 7.8 ± 0.2 |

### Benchmarks

Kilosort 2, 2.5, 3 and 4 outperformed all other algorithms in all cases. Kilosort 1 performed poorly, due to the lack of drift correction and its bias towards over-split units. The nearest competing algorithm in performance was IronClust developed by the Flatiron Institute, which accounts for drift in a different way from Kilosort [19]. IronClust generally found ∼ 50% of all units, compared to the 80-90% found by Kilosort4 (Table 2). Many of the algorithms tested did not have explicit drift correction. Some of these (SpyKING CIRCUS, MountainSort4) matched the IronClust performance at no drift, medium and fast drift, but their performance deteriorated drastically with higher drift [2, 3]. Among all algorithms with explicit drift correction (Kilosort 2.5, 3 and 4), Kilosort4 consistently performed better due to its improved clustering algorithm, and in some cases performed much better (on the step drift conditions). As we suspected, the aligned sites condition recovered the full performance of Kilosort4 on the step drift simulations, likely because it reduces the vertical sampling from 40 µm to 20 µm.

We also tested how well the drift amplitudes were identified by the drift detection algorithm from Kilosort2.5 (in the Kilosort4 implementation) and found good performance in all cases, except for the fast drift condition where the timescale of drift was faster than the 2 sec bin size used for drift correction (Figure S4). Much smaller bin sizes cannot be used for drift estimation, since a minimum number of spike samples is required. Nonetheless, the results show that Kilosort still performed well in this case, likely due to the robustness of the clustering algorithms. Finally, we calculated the performance of the algorithms as a function of the ground truth firing rates, amplitudes and spatial extents (Figure S5). The dependence of Kilosort4 on these variables was minimal. However, some of the other algorithms had a strong dependence on amplitude, which could not be improved by lowering spike detection thresholds. Also, many algorithms performed more poorly when the waveforms had a large spatial extent as opposed to having their electrical fields concentrated on just a few channels.

## Discussion

Here we described Kilosort, a computational framework for spike sorting electrophysiological data. The latest version, Kilosort4, represents our cumulative development efforts over the past eight years, containing algorithms like template deconvolution (from Kilosort1), drift correction (from Kilosort2.5), as well as completely new clustering algorithms based on graph methods. Furthermore, Kilosort4 was re-written from the ground up in Python, an open-source programming language, using the pytorch package for GPU acceleration. The popularity of pytorch/python should ensure that Kilosort continues to be further improved and developed. We have also developed a new simulation framework to improve the benchmarking of spike sorting algorithms. Our simulations contain realistic background noise and realistic drift with diverse properties, and they are qualitatively similar to real recordings with Neuropixels probes. Kilosort4 outperformed all other algorithms on all simulation conditions, in some cases by a large margin.

All versions of Kilosort have been developed primarily on Neuropixels data. However, since Kilosort adapts to the data statistics, it has been used widely on other types of probes and other recording methods. Some types of data do require special consideration. For example, some data cannot be drift corrected effectively due to either lacking a well-defined geometry (tetrodes), or due to the vertical spacing between electrodes being too high (more than 40 µm). This consideration also applies to data from single electrodes such as in a Utah array. Kilosort2 might be a better algorithm for such data, because it performs drift tracking without requiring an explicit channel geometry. Based on our benchmarks, Kilosort2 with drift tracking performs similarly to Kilosort2.5 with drift correction, except for the cases where step drift is present. Data from retinal arrays does not require drift correction and may be processed through Kilosort4, but it may require large amounts of GPU RAM for arrays with thousands of electrodes and thus would be better split into multiple sections and processed separately. Another special type of data are cerebellar neurons with complex spikes, which can have variable, complex shapes that are not well matched by a single template, and specialized algorithms for detection may be required [25]. Another special type of recording comes from chronic experiments over multiple days, potentially separated by long intervals. While we have not explicitly tested such recordings here, the benchmark results for the step drift simulation are encouraging because this simulation qualitatively matches changes we have seen chronically with implanted Neuropixels 2 electrodes [9].

The problem of identifying neurons from extracellular recordings has a long history in neuroscience. The substantial progress seen in the past several years stems from multiple simultaneous developments: engineering of better devices (Neuropixels and others), better algorithms (Kilosort and others), improved visualizations of spike sorting results (Phy) and multiple rounds of user feedback provided by a quickly-expanding community. Computational requirements have sometimes influenced the design of new probes, such as the aligned sites and reduced vertical spacing of Neuropixels 2 which were motivated by the need for better drift correction. Such computational considerations will hopefully continue to influence the development of future devices to increase the quality and quantity of neurons recovered by spike sorting.

## Acknowledgments

This research was funded by the Howard Hughes Medical Institute at the Janelia Research Campus.

## Author contributions

M.P. designed and built all versions of Kilosort. S.S. wrote the python GUI and C.S. developed the drifting simulations. C.S. and M.P. performed data analysis, coordinated the project and wrote the paper.

## Code availability

Kilosort4 will be available upon publication at https://www.github.com/mouseland/kilosort. Version 2, 2.5 and 3 are currently available at the same link.

## Data availability

We used datasets shared by Nick Steinmetz and the International Brain Laboratory [13, 15]. The datasets are available at at http://data.cortexlab.net/singlePhase3/ and https://ibl.flatironinstitute.org/public/.

## Methods

The Kilosort4 code library is implemented in Python 3 [10] using pytorch, numpy, scipy, scikit-learn, faiss-cpu, numba and tqdm [11, 26–32]. The graphical user interface additionally uses PyQt and pyqtgraph [12, 33]. The figures were made using matplotlib and jupyter-notebook [34, 35]. Kilosort 2, 2.5 and 3 were implemented in MATLAB.

We demonstrate the Kilosort4 method step-by-step in Figure 1 and Figure 2. In Figure 1 an electrophysiological recording from Nick Steinmetz was used (“Single Phase 3”; [13] and https://figshare.com/articles/_Single_Phase3_Neuropixels_Dataset/7666892). In Figure 2 an electrophysiological recording from the International Brain Laboratory was used (id: 6f6d2c8e-28be-49f4-ae4d-06be2d3148c1; [15]). Both recordings were performed with a Neuropixels 1.0 probe, which has 384 sites organized in rows of two with a vertical spacing of 20 µm, a horizontal spacing of 32 µm. Due to the staggered design (16 µm horizontal offset between consecutive rows), the spatial repetition period of this probe is 40 µm.

### Graphical user interface (GUI)

We developed a graphical user interface to facilitate the user interaction with Kilosort4. This interface was built using pyqtgraph which itself uses PyQt [12, 33], and it replicates the Matlab GUI which was originally built for Kilosort2 by Nick Steinmetz. The GUI allows the user to select a data file, a configuration file for the probe, and set the most important parameters manually. In addition, a probe file can be constructed directly in the GUI. After loading the data and configuration file, the GUI displays a short segment of the data, which can be used to determine if the configuration was correct. Typical mistakes are easy to identify. For example if the total number of channels is incorrect, then the data will appear to be diagonally “streaked” because multi-channel patterns will be offset by 1 or 2 extra samples on each consecutive channel. Another typical problem is having an incorrect order of channels, in which case the user will see clear single-channel but no multi-channel waveforms. Finally, the GUI can produce several plots during runs which can be used to diagnose drift correction and the overall spike rates of the recording.

### Algorithms for Kilosort4

In the next several sections we describe the algorithmic steps in Kilosort4. Some of these steps are inherited or evolved from previous versions. For clarity, we describe each of the steps exactly as they are currently used in Kilosort4. If a previous version of Kilosort is different, we clearly indicate the difference. We dedicate a completely separate section below for algorithms not used in Kilosort4 but used in previous versions.

Many of the processing operations are performed on a per-batch basis. The default batch size is *N*_*T*_ = 60, 000, and it was *N*_*T*_ = 65, 536 in versions 2/2.5/3 and *N*_*T*_ = 32, 768 in version 1. The increase in batch size in Kilosort2 was designed to allow better per-batch estimation of drift properties. Due to the per-batch application of temporal operations, we require special considerations at batch boundaries. Every batch of data is loaded with left and right padding of *n*_*t*_ additional timepoints on each side (*n*_*t*_ = 61 by default). On the first batch, the left pad consists of the first data sample repeated *n*_*t*_ times. The last batch is typically less than a full batch size of *N*_*T*_. For consistency, we pad this batch to the full *N*_*T*_ size using the repeated last value in the data.

The clustering in Kilosort 3/4 is done in small 40 µm sections of the probe, but including information from nearby channels and including spikes extracted at all timepoints.

### Preprocessing

Our standard preprocessing pipeline includes a sequence of operations: common average referencing (CAR), temporal filtering, channel whitening and drift correction. In Kilosort4, all these steps are performed on demand whenever a batch of data is needed. In all previous versions, the preprocessing of the entire data was done first and the preprocessed data was stored in a separate binary file. Drift correction was introduced in Kilosort 2.5.

#### Common average referencing

The first operations applied to data are to remove the mean across time for each batch, followed by removing the median across channels (common average referencing or CAR). The CAR can substantially reduce the impact of artifacts coming from remote sources such as room noise or optogenetics. The CAR must be applied before the other filtering and whitening operations, so that large artifacts do not “leak” into other data samples.

#### Temporal filtering

This is a per-channel filtering operation which defaults to a high-pass filter at 300Hz. Bandpass filtering is typically done using IIR filters for example with Butterworth coefficients. Butterworth filters have some desirable properties in the frequency space, but their implementation on the GPU is slow. To accelerate it, we switch to using an FIR filter that simulates the Butterworth filter and we perform the FIR operation in FFT space taking advantage of the convolution theorem. To get the impulse response of a Butterworth filter, we simply filter a vector of size *N*_*T*_ with all zeros and a single 1 value at position floor(*N*_*T*_ */*2) (0-indexed).

#### Channel whitening

While temporal filtering reduces time-lagged correlations coming from background electrical activity, it does not reduce across-channel correlations. To reduce the impact of local sources, such as spikes from 100-1000µ*m* away from the probe, we perform channel whitening in local neighborhoods of channels. A separate whitening vector is estimated for each channel based on its nearest 32 channels using the so-called ZCA transform, which stands for Zero Phase Component Analysis [36]. ZCA is the data whitening transformation which is closest in Euclidean norm to the original data. For an *N* by *T* matrix *A*, the ZCA transform matrix *W* is found by inverting the covariance matrix, using epsilon-smoothing of the singular values:

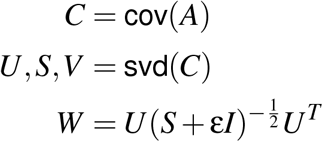

The local whitening matrix *W* is calculated separately for each channel and its neighborhood of 32 channels, and only the whitening vector corresponding to that channel is kept and embedded into a full-size *N*_*chan*_ by *N*_*chan*_ matrix. This is preferable to directly calculating a grand *N*_*chan*_ by *N*_*chan*_ whitening matrix because it reduces the number of whitening coefficients to 32 · *N*_*chan*_ instead of *N*_*chan*_ · *N*_*chan*_ which prevents overfitting in the limit of a large *N*_*chan*_.

#### Drift correction

Drift correction is a complex preprocessing step which was described in detail in [9]. Here we only described a few small modifications in Kilosort4. The drift correction process can be separated into drift estimation and data alignment. In Kilosort4, drift estimation is performed in advance, while data alignment is performed on-demand along with the other preprocessing operations. Drift estimation includes a step of spike detection, which uses a set of predefined, “universal” templates to detect multi-channel spikes. In Kilosort 2.5 and 3, these predefined templates were constrained to be negative-going spikes, while in Kilosort4 we consider both positive and negative going spikes using pairs of inverted templates (for fast computation). Another modification in Kilosort4 is the use of linear interpolation for sampling the drift traces at every channel, in place of the “Makima” method used in previous versions.

Since data alignment is a linear operation performed with a Gaussian kriging kernel, it can be combined with channel whitening which is also a linear operation. In practical terms, the two *N*_*chan*_ by *N*_*chan*_ matrix multiplications are combined into one, thus further accelerating the computation.

### Template deconvolution

Template deconvolution is the process of using a set of waveform templates matched to the data in order to detect spikes and extract their features, even when they overlap other spikes on the same channels and at the same timepoints. Template deconvolution can be seen as replacing the spike detection step in a classical spike sorting pipeline. The goal in Kilosort4 is to extract all the spikes above a certain waveform norm, and calculate their spike features in a way that discards the contribution of nearby overlapping spikes. Template deconvolution improves on classical spike detection in several ways:

1. The detection of the spikes is performed by template matching, which is a more effective way of detecting spikes compared to threshold crossings, because it uses templates that represent the multi-channel spikes of the neurons being matched.
2. Spikes that overlap in time and channels can be detected and extracted as separate events due to the use of iterative matching pursuit. Classical methods require an “interdiction” area in time and channels around each detected spike where a second spike detection is disallowed, in order to prevent double detections of the same spike.
3. The features extracted for each spike can be de-contaminated from other overlapping spikes, due to the use of a generative or reconstructive model. As described below, these features are robust to imperfectly chosen templates.

#### Template learning

To perform template deconvolution, a set of templates must be learned that can match all the detectable spikes on the probe. In previous Kilosort versions (1 / 2 / 2.5), special care was taken to ensure that these templates match neural waveforms on a one-to-one basis. This was necessary because relatively few additional merges and splits were performed after template deconvolution. In Kilosort 3 and 4, the templates do not need to match single neurons because the features extracted by template deconvolution are clustered again using more refined clustering algorithms. However, it is important that every spike in the raw data has *some* template to match to.

To build a set of templates, we perform clustering on a set of spikes identified by template matching with a set of universal spike templates. This initial spike detection step is equivalent to the spike detection performed in Kilosort 2.5 for drift correction. The universal templates are defined by all possible combinations of a spatial position in 2D; 2) a single-channel waveform shape; 3) a spatial size. The spatial positions need not be coincident with actual probe channels, and we choose them to upsample the channel densities by a factor of 2 in each dimension. For a Neuropixels 1 probe, this corresponds to 1536 positions. The single-channel waveform shapes are obtained by k-means clustering of single channel spikes, either from a pre-existing dataset (IBL dataset) or from spikes detected by threshold crossings in the data, and we default to 6 such waveforms. Finally, the spatial sizes (five by default) define the envelope of an isotropic Gaussian centered on the spatial position of the template, which is used as per-channel amplitudes. In total, a set of 46,080 universal templates are used for a Neuropixels 1 probe; for more details see [9]. The spatial footprints are explicitly precomputed for all positions and all spatial sizes. The templates are effectively normalized to unit norm by separately normalizing the per-channel waveform templates and the spatial footprints. Since the universal templates are unit norm, their variance explained at each timepoint can be easily calculated as the dot product with the data, squared:

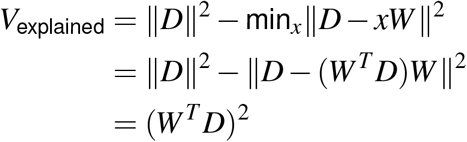

where *W* is the unit-norm universal template, *D* is the data over a particular set of channels and time-points, and *x* is the best matching amplitude that the template needs to be multiplied by to match the data.

The dot products between each of these templates and the data at each timepoint can be performed efficiently in the following order: 1) temporal convolution of each data channel with each of the 6 single-channel waveforms; 2) per timepoint matrix multiplication with a set of weights corresponding to all positions and all spatial sizes. Once the dot products are calculated in this manner, the largest variance explained value is kept at each spatial position of each template. For a Neuropixels probe, this is a matrix of size 1536 by *N*_*T*_ (batch size). The goal of this spike detection step is to find localized peaks in this matrix, which must be local maxima in a neighborhood of timepoints (± 20) and spatial positions (100 nearest positions). The relatively large neighborhood size ensures that no spike is detected twice, but prevents many overlapping spikes from being detected (typically about 50% of spikes go undetected). However, the missing spikes are not a concern for the purpose of template learning, since it is extremely unlikely that all the spikes from a neuron will be consistently missed by this procedure.

Once the spikes are detected, we extract PC features in the 10 nearest channels to each detection. We use a set of six PCs which are found either from a pre-existing dataset (IBL dataset) or from spikes detected by threshold crossings. For each spike, an XY position on the probe is computed based on the center-of-mass across channels of the spike’s projection on the best-matching single channel template (same as in Kilosort 2.5). We assign all spikes in 40 µm bins according to their vertical position, and embed all spikes detected in the same bin to the same set of channels (which is usually more than 10 channels due to differences between spike positions). Finally, the embedded PC features are clustered according to the same graph-based clustering algorithm we describe below, using only the merging criterion of the bimodal regression-axis and not using the cross-correlation based criterion. In Kilosort3, the same procedure is applied but the clustering algorithm is recursive pursuit. After clustering each 40 µm section of the probe, the centroids are multiplied back from PC space into spatio-temporal waveforms, and pooled together across the probe.

Templates from the same neuron may be detected multiple times, either on the same 40 µm section or in nearby sections. This is not inherently a problem because each neuron can have multiple templates. However, it can become a problem if these multiple templates are not aligned to each other, because then spikes from the same neuron will be detected at different temporal positions, which changes their PC feature distribution. In addition, having many templates makes the spike detection step memory and compute inefficient. A solution to both these problems is to merge together templates which have a high correlation with each other and similar means, where the correlation is maximized across possible timelags. In addition, we temporally align all templates based on their maximal correlation with the same six prototypical single-channel waveforms describe above. Note that this merging step may result in the opposite scenario of having one template for multiple neurons. This is also not a problem, because templates are only merged when they have a high correlation, and thus the same average template can successfully match the shape of multiple neurons.

#### Spike detection with learned templates and matching pursuit

Once a set of templates is learned, they can be used for template matching similar to the universal templates described above. The main difference is that instead of allowing for an arbitrary scaling factor *x*, we require that matches use the average amplitude of the template it was found with. The variance explained of learned template *W* of some data *D* thus becomes:

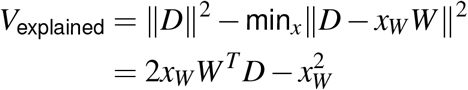

Like before, this quantity only requires the calculation of *W*^*T*^ *D*, which can be done convolutionally for each template. In practice, we represent templates using a three-rank approximation, factorized over channels and time, which speeds up the convolutions dramatically [6]. We first multiply the data with the channel weights for each rank, and convolve the resulting traces with the temporal components. The three-rank approximation captures nearly the entire waveform variance in all cases ([6]), and also helps to denoise templates calculated from relatively few spikes.

To extract overlapping spikes, we must detect spikes iteratively over the same portion of data, and subtract off from the data those parts attributed to spike detections. This subtraction allows for another pass of detections to be performed, which can detect other spikes left over and yet un-subtracted. This procedure is called matching pursuit ([37]) and is fundamentally a sequential process: to detect another spike, one must first subtract off the contributions of spikes detected before. However, we can parallelize this step thus making it suitable for GPU processing by observing that the subtraction of a single spike results in highly-localized changes to the data, which cannot affect the calculated amplitudes far from the position of that subtracted spike. Thus, we can detect and subtract multiple spikes in one round as long as they are far enough from each other. Upon calculating a matrix of variance explained for each template at each timepoint, we detect peaks in this matrix which are local maxima over local neighborhoods in time ± *n*_*t*_ time samples, and across all channels. After detection, the optimal amplitude for each spike is calculated and its contribution from the data is subtracted off. To avoid recalculating the dot products of templates at all timepoints, the contribution of the subtracted spikes to the dot-products is directly updated locally using a set of precomputed dot-products between templates, at all possible timelags. This detection and subtraction process is repeated for 50 rounds, with later rounds being much faster due to the increasingly smaller number of spikes left to extract.

#### Extracting PC features with background subtraction

The final step in template deconvolution is to extract features from the data to be used by the clustering algorithm. One possibility would be to directly extract PC features from the preprocessed data, at the spike detection times (Figure 1h), however this results in contamination with background spikes. A better option is to first subtract the effect of other spikes, since we know from the matching pursuit step how much these other spikes contribute (Figure 1e). To do this computation efficiently, we first extract PC features from the residual (Figure 1f), and then add back to these features the contribution of the template which was used to extract the spike. The contribution of each template in PC space is precomputed for faster processing.

### Graph-based clustering

The new clustering algorithm in Kilosort4 uses graph-based algorithms. This class of algorithms relies entirely on the graph constructed by finding the nearest neighbors to each data point. There are several steps:

1. Neighbor finding with subsampling
2. Iterative neighbor reassignment
3. Hierarchical linkage tree

#### Neighbor finding with subsampling

Many frameworks for fast neighbor finding exist and we tested a lot of them for spike sorting data. In the end, the brute force implementation from the faiss framework [38] outperformed other approaches in speed on modern multi-core computers for the range of data points we need to search over (10,000-100,000) and the number of data points we need to find neighbors for (100,000-1,000,000).

#### Iterative neighbor assignment

Clustering algorithms based on graphs typically optimize a cost function such as the modularity cost function. We review this approach first, before describing our new approach. Following [17], the modularity cost function is defined by

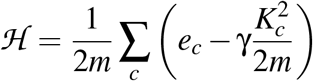

where *m* is the total number of edges in the graph, *e*_*c*_ is the number of edges in community *c, K*_*c*_ is the sum of degrees in community *c* and γ is a “resolution” parameter that controls the number of clusters. The 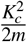 can be interpreted as the expected number of edges in community *c* from a null model with the same node degrees as the data but otherwise random graph connections.

Specialized optimization algorithms exist to maximize the modularity cost function by moving nodes between communities and performing merges when the node re-assignment converges [18]. Additionally, splitting steps and other optimizations were recently introduced which improve the results of the algorithm and its speed [17]. These algorithms are effective for many types of data, yet have a substantial failure mode for spike sorting data: they have difficulty clustering data with very different number of points per cluster. In practice, for our clustering problems, there are often very large clusters of up to 100,000 points together with clusters with many fewer (*<*1,000) points. A low resolution parameter γ can keep the large cluster in one piece, but also merges the small clusters into larger clusters. Conversely, high resolution parameters may return the small clusters as individual clusters, but can split the large cluster into very many (hundreds) of pieces. The oversplitting is not inherently a bad property, since we will perform merges on these clusters anyway, but the very large number of pieces returned for the large clusters means that very many correct merging decisions must be made, which is in itself a very difficult optimization problem. In addition, running the Louvain/Leiden algorithms with large resolution parameters may somewhat reduce the effectiveness of the algorithm since the community penalty 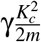 only has a null model interpretation for λ = 1.

To improve on these algorithms, we started from the observation that local minima of the neighbor re-assignment step have some desirable properties. These local minima arise because the neighbor reassignment step monotonically improves the modularity cost function by greedily moving nodes to new clusters if that improves the modularity score. This step converges after a while, because no more clusters can be moved. This is however a local minimum of the optimization, and the modularity can often be further increased by making merges between clusters. Unlike the node re-assignment, which consists of small local moves, the merging between clusters is a global move in the cost function and can thus escape the local minimum. Algorithms like Leiden/Louvain take advantage of such global merges by applying the node re-assignment step again on a new graph made by aggregating all the points into their clusters when the local minimum is reached.

Our observation was that the local minima themselves can consist of good clustering, if the neighbor re-assignment step is initialized appropriately. Our initialization uses the K-means++ algorithm to partition the data initially into 200 clusters [14]. The node reassignment algorithm for the modularity cost function with γ = 1 is run for a fixed number of iterations (typically sufficient for convergence). The converged partitioning of the data is then used as a clustering result. Especially relevant to the next step, the algorithm almost never made incorrect merges, and instead output some clusters oversplit. This bias towards oversplitting is important, because it allows us to correct the mistakes of the algorithm by making correct merge decisions, which is much easier than finding the correct split in a cluster.

We also found that clusters which were oversplit generally had a reason to be oversplit: the separate pieces identified by the algorithm were in fact sufficiently different to create a local minimum in the cluster assignments. This is a common problem in spike sorting data, where nonlinear changes in the waveform can result in clusters that appear bimodal in Euclidian space. An extreme example of this effect is due to abrupt drifts of the probe changing the sampling of the waveforms by a non-integer multiple of the probe period. Even after drift correction, waveforms sampled at the two different positions will be much more similar to other waveforms from the same position, than they are to waveforms sampled at the other position (Figure S2b). As a consequence, many algorithms return such units oversplit into two halves, as can be clearly seen in the benchmark results for the step drift condition, where many units are identified with exactly a 0.5 score, which corresponds to 50% of the spikes identified.

#### Hierarchical merging tree

To perform merges, we could take two strategies: 1) a brute-force approach in which we check all pairs of clusters for merges, or at least the ones with high waveform correlation; 2) a directed approach where we use the structure of the data to tell us which merges to check. We use both, starting with the second one to reduce the number of clusters and thus reduce the number of brute-force checks we need to make later.

For the directed approach, we construct a hierarchical merging tree based on the modularity cost function. The leaves of this tree consist of the clusters identified at the previous step. For each pair of clusters *i, j*, we aggregate the neighbors and node degrees, similar to the Leiden/Louvain algorithms, thus resulting in a full matrix *K* of size *n*_*k*_ by *n*_*k*_ where *n*_*k*_ is the number of clusters, and where *K*_*i j*_ is the number of edges between clusters *i, j*, while *K*_*ii*_ is the number of internal edges. Additionally, a variable *k*_*i*_ holds the aggregated degree of each cluster *i*. The linkage tree is constructed by varying the resolution parameter γ in the modularity cost function from ∞ down to 0. As γ decreases, merges of two clusters start to increase the modularity cost function. Specifically, a pair of clusters gets merged when the modularity ℋ_2_ after merging equals the modularity ℋ_1_ before merging, where:

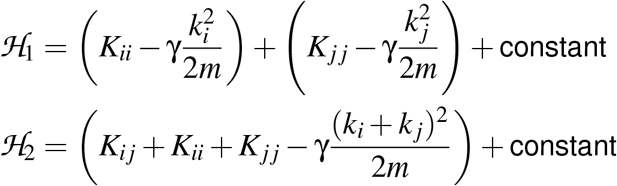

Setting ℋ_2_ = ℋ_1_ yields:

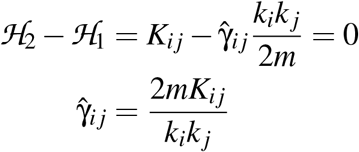

In other words, a pair of clusters *i, j* should be merged when γ reaches a value of 2*mK*_*i j*_*/*(*k*_*i*_*k*_*j*_). After merging, the matrix *K* and vector *k* can be recomputed with the two clusters *i, j* becoming aggregated into one. Note that a merging decision does not change the 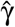 for other pairs of clusters, and it cannot result in a higher 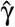 than the current 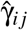. This can be shown by reductio ad absurdum: if the merged *i, j* cluster had a higher 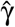 with another cluster *l*, it would imply that one of the original clusters *i* or *j* had a higher 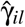 or 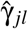 _*jl*_, and thus it should have been merged a priori. The mono-tonic property of 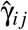 ensures that a well-defined merging tree exists, with a strictly decreasing sequence of 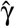 for increasingly higher merges in the tree. Empirically, we have found that the resulting merging tree is very useful for making merge/split decisions.

### Split/merge criteria

With the tree constructed, we next move down the tree starting from the top and make individual merge/split decisions at every node. If a node is not being split, then the splits below that node are no longer checked. We use two splitting criteria: 1) the bimodality of the data projection along the regression axis between the two clusters and 2) the degree of refractoriness of the cross-correlogram. If the pair of units has a refractory cross-correlogram, then the split is always performed. If the cross-correlogram is not refractory, then the split is performed if and only if the projection along the regression axis is bimodal.

#### Bimodality of regression axis

Consider a set of spike features **x**_*k*_ with associated labels *y*_*k*_ ∈ {−1, 1}, where − 1 indicates the first cluster and 1 indicates the second cluster. A regression axis 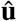 can be obtained by minimizing:

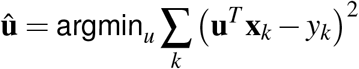

This regression problem becomes highly unbalanced when one of the clusters has many more points than the other. We therefore add a set of weights *w*_−1_ = *n*_2_*/*(*n*_1_ + *n*_2_), *w*_+1_ = *n*_1_*/*(*n*_1_ + *n*_2_), where *n*_1_, *n*_2_ are the number of spikes in the first and second cluster.

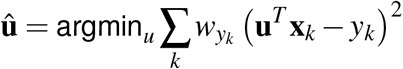

This weighted regression problem can be solved in the usual fashion. Finally, we use the 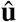 axis to estimate how well separated the clusters are by projecting 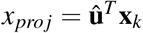. The projections are binned in 400 bins linearly-spaced between -2 and 2, and the histogram is gaussian smoothed with a standard deviation of 4 bins. To score the degree of bimodality, we find three important values in the histogram: the peak of the negative portion, the trough around 0, and the peak of the positive portion. First we find the trough *x*_*min*_ at position *i*_*min*_ in the bin range of 175 to 225. Then we find the peaks *x*_1_, *x*_2_ in the bin ranges from 0 to *i*_*min*_ and from *i*_*min*_ to 400. The bimodality score is defined by

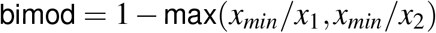

In other words, we compare the density of the *x*_*pro j*_ distribution at its trough to the peak densities for both clusters. If the density at the trough is similar in value to the density of either the left or right peak, that indicates a non-bimodal distribution.

#### Refractory auto- and cross-correlograms

There are many cases where the regression axis has a bimodal distribution, yet the clusters are part of the same neuron. This is due to the non-stationarity of the waveforms from the same neuron, either due to drift or due to other factors. In such cases, we need to use extra information such as the statistics of the spike trains. Fortunately, all neurons have a refractory period, which is a short duration (1-5ms) after they fire an action potential when they cannot fire again. The refractory period is heavily used by human curators to decide whether: 1) a cluster is well isolated and not contaminated with spikes from other neurons; 2) a pair of clusters are distinct neurons or pieces of the same neuron. These two decisions can be made based on the auto- and cross-correlograms (ACG and CCG) respectively:

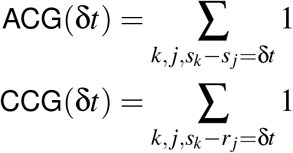

where *s*_*k*_, *r*_*j*_ represent the spikes times of the two neurons. In practice, we bin the auto- and cross-correlograms in 1ms bins from δ*t* = −0.5 sec to δ*t* = 0.5 sec. We consider the central bins of the cross-correlograms, and calculate how likely it is to see a very small number of coincidences in that bin, if the two clusters are from neurons firing independently from each other. We define *n*_*k*_ as the number of coincidences in the central − *k* to +*k* bin range, *R* as the baseline rate of coincidences calculated from the other bins of the cross-correlogram. Cross-correlograms may be assymetric, and to account for that we estimate *R* as the maximum rate from either the left or right shoulder of the cross-correlogram. We use two criteria to determine refractoriness. The first criterion is simply based on the ratio of refractory coincidences versus coincidences in other bins which works well in most cases, except when one of the units has very few spikes, in which case very few refractory coicindences may be observed just by chance. For the first criterion, we use the ratio *R*_12_ of *n*_*k*_ to its expected value from a rate *R*, where *R*_12_ takes the minimum value of this ratio across *k*. We set a threshold of 0.25 on *R*_12_ to consider a CCG refractory, and 0.1 to consider an ACG refractory. For the second criterion, we use the probability *p*_*k*_ that *n*_*k*_ spikes or less would be observed from a Poisson process with rate λ_*k*_ = (2*k* + 1)*R*, which we approximate using a Gaussian with the same mean and standard deviation as the Poisson process as

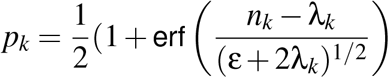

where ε = 10^−10^ is a small constant to prevent taking the square root of 0. If *Q*_12_ = min(*p*_*k*_) is large, it implies that the number of refractory spikes have a high chance of being observed from a Poisson distribution with the baseline rate, and thus the CCG is not refractory. We set a threshold on *Q*_12_ of 0.05 to consider a CCG refractory, and 0.2 to consider an ACG refractory. Both criteria have to be satisfied for a CCG to be refractory: *R*_12_ *<* 0.25 and *Q*_12_ *<* 0.05 for the CCG and *R*_12_ *<* 0.1 and *Q*_12_ *<* 0.2 for the CCG. The different thresholds for ACG and CCG has to do with the function of these decisions: for the ACG, we want small contamination rates *R*_12_ because it indicates a well-isolated neuron, while for the CCG we want to prevent clusters from being split if their contamination rate *R*_12_ is indicative of a relation between these two clusters. Similarly for *Q*_12_.

#### Global merges

Global merges are performed after all sections of the probe have been clustered. As a similarity metric, we use the maximum correlation of pairs of waveforms over all timelags. To test for merges, we sort all units by their number of spikes, and start testing in order from the units with the most spikes. For each unit, we find all other units with a similarity above 0.5 and start testing for merges starting from high to low similarity. A merge is performed if the cross-correlogram is refractory. After a merge is performed, the merged unit is retested again versus all other units with similarity above 0.5 with it. After no more merges can be performed, a unit is considered “complete”, and is removed from potential merges with subsequent tested units.

### Scaling up the graph-based clustering

Graph-based clustering algorithms do not scale well with the number of data points, and we had to develop new formulations and optimization strategies. The poor scalability is due to several problems: 1) finding the neighbors of all points scales quadratically with the number of points; 2) the *K* nearest neighbors in a small dataset are relatively further away from the *K* nearest neighbors in a larger dataset; 3) existing optimization algorithms like Leiden/Louvain are inherently sequential and thus hard or impossible to parallelize on GPUs. The first problem could be reduced by using some of the neighbor finding algorithms that have sublinear time for finding neighbors [27]. However, for the particular type of data we consider, we find these algorithms to be slower, not faster than the brute force approach, at least when a multi-core CPU is used. The second problem is an issue because the effective neighborhood size around a point influences its clustering properties. If the neighborhood sizes are very small, clusters may split up into multiple pieces more easily. If it is too large, it may include points from other clusters. As a recording grows in duration, the number of spikes grows linearly with it. Thus, some normalization step must be introduced to ensure that neighborhood sizes are comparable for short and long recordings. To solve the third problem, a redesign of the cost function is necessary, so as to make multiple optimization steps in parallel.

Our approach for improving scalability relies on a subsampled data approach, where we only search for neighbors in a smaller subset of all points. In other words, instead of constructing an *N* by *N* adjacency matrix, where *N* is the number of points, we construct an *N* by *n*_*sub*_ adjacency matrix, where *n*_*sub*_ is a fixed number of spikes independent of recording length, which is determined by the size of the section of the probe being clustered (40 µm typically, for which we use *n*_*sub*_ = 25,000). This solves the first two problems, but not the third. To solve the third problem, we treat the adjacency graph as a bipartite graph, by designating the subsampled datapoints as a different set of nodes, which we will call the “right” nodes, as opposed to the “left” nodes which consist of all datapoints. Note that this is purely a mathematical construction, in which we duplicated the subsampled nodes so they exist both among the left and the right nodes. The reason for making the graph bipartite is to allow the cluster identities for left nodes to be optimized independently, given the identities of the right nodes, and viceversa. However, making the graph bi-partite is not sufficient, we must also modify the modularity cost function from:

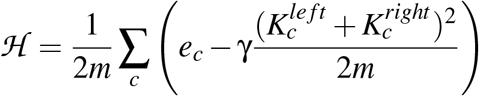

into:

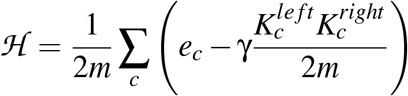

where 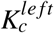 is the sum of degrees of left nodes in the cluster *c*, 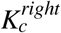 is the sum of degrees of right nodes, and *e*_*c*_ are the number of edges between left and right nodes. If the cluster identities for all right nodes are fixed, a short calculation shows that every left node *t* can be assigned independently to a cluster σ_*t*_ to maximize their contribution to the modularity cost function:

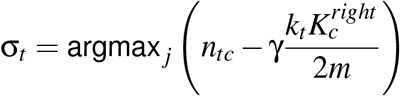

where *n*_*tc*_ are the number of right node neighbors of left node *t* in cluster *c*, and *k*_*t*_ is the degree of node *t* like before. Similarly, every right node can be assigned independently given fixed assignments for all left nodes. Thus, we can iterate between assigning cluster identities to all right nodes given all the left nodes, followed by assigning all the left nodes given all the right nodes. Note that a left node which represents the same point as a right node may in fact be assigned to a different cluster than its corresponding right node. This new iterative optimization has massive parallelism, and thus is suitable for GPU acceleration.

This optimization is initialized with 200 clusters identified by K-means++, which we implemented in pytorch for GPU-based scalability [14].

### Algorithms for Kilosort 2/2.5/3

The previous section completes the description of Kilosort4. In the next sections, we decribe algorithms from previous versions of Kilosort which have not been previously described. These are divided into drift tracking (Kilosort2), global optimization (Kilosort2/2.5) and recursive pursuit (Kilosort3).

### Drift tracking (Kilosort 2)

Drift tracking was an alternative strategy of accounting for drift. Unlike the drift correction algorithms from Kilosort2.5 and onwards, drift tracking does not require a geometrical model of the recording channels and thus it can be used for recordings with tetrodes, single electrodes etc. Drift tracking works well when drift changes are continuous, or at least the drift positions overall span a continuous range. Drift tracking does not work well when the recording consists mainly of two drift positions, with little sampling of the positions in-between (see step drift benchmarks in Figure 3). Drift tracking requires two algorithmic steps, described below: online template learning and fast drift tracking.

#### Online template learning and tracking

In the simplest case, imagine that the drift of the probe is very slow. A possible spike sorting strategy in that case could be to start by spike sorting a subsection of the data, say 5 minutes, over which the probe is at an almost fixed position, since drift is slow. With the templates learned from 5 minutes of data, one could then use the templates to extract and assign spikes on the next 5 minutes of data, and update the templates based on the spikes that were found. If the drift is slow, the distribution of waveforms from single templates will have only shifted slightly, so that spike assignments to clusters are still correct. The update of the templates would then track the mean of the shifted distribution for each cluster. This is, in a broad sense, the drift tracking strategy from Kilosort2, and it was a natural extension of the online template learning of Kilosort1. In the rest of this section, we describe the exact mathematical form of online template learning.

The generative model of Kilosort is given by the reconstruction cost function:

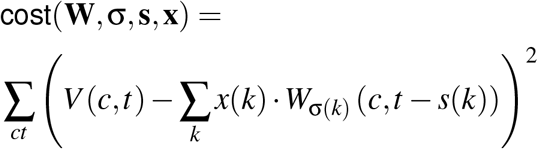

where *V* (*c,t*) is the recorded voltage at channel *c* and timepoint *t*, σ(*k*) is the template index for spike *k* at time *s*(*k*), *W*_*i*_ is the multi-channel template for cluster *i* and *x*(*k*) is the amplitude of spike *k*. For mathematical simplicity, we assume “infinite” temporal windows for each template *W*_*i*_ and we also assume that they span all channels of the probe, but in practice we restrict each template to a width of *n*_*t*_ = 61 samples and to a small number of channels (typically 32). Learning and inference in this model proceeds via the standard “EM-style” algorithm. For inference, we assume the templates *W* are fixed, and we simultaneously infer *s*(*k*), σ(*k*), *x*(*k*) for all *k* via the parallelized matching pursuit algorithm described above. Learning proceeds iteratively by inferring **s**, σ, **x** from a single batch with the current *W*, and computing an improved *W* for this batch:

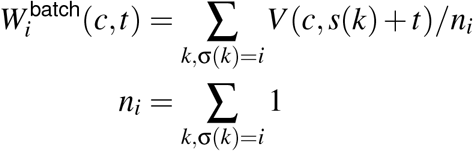

where we omit the dependence on amplitudes *x*(*k*) for robustness, since outlier artifacts may have very large *x*_*k*_ that could dominate the templates. To convert this EM-style algorithm into an online algorithm, we perform the inference at a single batch level (≈ 2 sec), and update the templates with an exponential filter which depends on the number of spikes inferred for each template:

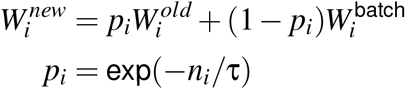

where τ is typically set to 400 spikes. In other words, it takes approximately 400 new inferred spikes for *W*_*i*_ to “forget” its previous value. For learning to be effective, the batches from one recording are processed in pseudorandom order in Kilosort1. For tracking in Kilosort2, we fix the order of the batches. For example, if the batches were processed in consecutive order 1, 2, 3, etc, then online learning of the templates would ensure tracking of the slow changes in templates over long timescales, similar to the simple scenario describe at the beginning of this section. In practice however, drift often contains fast components in addition to slow components. Since the τ scale is set to 400 spikes, fast drift cannot be tracked well with this approach. Reducing τ would improve tracking, at least for neurons with high firing rates, but many neurons fire at ∼ 1Hz and/or in bursty sequeces. For such neurons, drift movements *<*1 min would be quite difficult to track, since very few spike samples of the neurons are seen in that time.

#### Tracking fast drift

To track fast drift, we make another modification to the online template learning algorithm. Instead of processing the batches in random order (like in Kilosort1), or in consecutive order (like for slow drift), we use a special re-ordering of the batches which puts similar batches next to each other. We define and estimate a drift dissimilarity metric between batches, based on the distributions of spike shapes in each batch.

To construct the drift similarity metric, the first step is to extract a set of templates **W**^*k*^ for each batch *k*. These templates are obtained by first detecting spikes via threshold crossing of PC-projected data. The PC projection is performed by convolving the data with the top three PCs, squaring and adding together the projections. Local maxima in this projection are found in a neighborhood of the nearest 17 channels and 61 timepoints. The features are extracted at the local maxima via PC projection for a subset of neighboring channels. Scaled k-means clustering is then performed, which is initialized with a random subset of the spikes and implemented on the GPU for speed, since it needs to be performed once for each 2 sec batch. A fixed number of clusters is used, equal to half the total number of channels. The centroids of the clustering are used as templates.

Once the templates **W**^*k*^ are obtained for each batch, we calculate a dissimilarity matrix *A*_*i j*_ between each pair of batches, each with their own template sets *W*^*i*^, *W* ^*j*^:

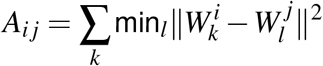

In other words, *A*_*i j*_ is a metric which is small when every template *k* in a set of templates *W*^*i*^ has a close match in another set of templates *W* ^*j*^. Pairs of batches *i, j* from similar drift levels will therefore have a small dissimilarity, while batches taken at very drift levels will have high dissimilarity because the templates won’t be very well matched.

Once we compute the matrix **A**, all that is left is to find a permutation ρ of the batches in which small dissimilarity values *A*_ρ(*i*)ρ(*j*)_ are near the diagonal. The algorithm we use for this is a version of “rastermap”, a framework algorithm we have been developing for sorting high-dimensional data along a one-dimensional continuum. This particular version of rastermap matches the similarity matrix **A** to the matrix of distances in a one-dimensional space, where *x*_*k*_ is a scalar value assigned to each batch *k* and optimized by the algorithm:

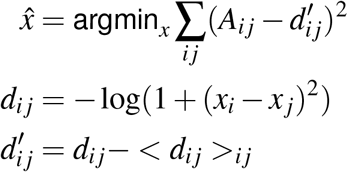

where the *<*·*>* operation signifies averaging. This cost function is initialized with *x*_*k*_ based on the largest left singular vectors of **A** and minimized by gradient descent. We preprocess **A** by z-scoring each row separately and symmetrize it by adding its transpose to it. In the optimization we ignore the mean of *d*_*i j*_ over all *i, j*, to avoid having to fit a constant offset term. Once the gradient descent optimization converges, we obtain a sorting of the batches by ρ = argsort(**x**), where the argsort operation returns the index order of a vector.

This ordering ρ is used to perform online template learning and tracking. The template learning is performed by running the algorithm over one half of the data (typically the first half after reordering, from the middle of the ρ range to the first batch and then back to the middle of the range). During this stage, the templates are being learnt: new templates can be introduced from the residuals of the reconstruction process, templates which are not used above a baseline spike rate are discarded, and merges and splits are also performed. See the next section for this template learning step. Once the template set is learned, the tracking is performed from the middle of the ρ range to the first batch, and then from the middle of the ρ range to the last batch. During the tracking phase, no new templates can be created or destroyed with any of the operations performed in the template learning phase. However, the template waveform itself continues to change in an online fashion as described above in the online template learning section.

### Global optimization algorithms for clustering (Kilosort 2 / 2.5)

Drift tracking was not the only new algorithm introduced in Kilosort2. We also added algorithmic steps designed to perform global optimization moves, in order to escape local minima which are very common in clustering algorithms. These global optimization moves were of three types: initialization, splitting a cluster and merging two clusters. We describe these in separate sections below. Some global optimization moves were also performed in Kilosort1 (simple splits and merges), but they were not as important there because the automated results of Kilosort1 underwent manual curation in Phy, and thus an oversplit cluster distribution was preferred because merges are much easier than splits. For Kilosort2 however, the automated results of the algorithm became sufficiently good to be used in an automated manner, and thus it was important to avoid oversplit or overmerged clusters.

#### Template initialization from residual

Initialization is one of the most important steps in a clustering algorithm. A common initialization for k-means style algorithms is k-means++, which sequentially adds data points as cluster centroids if they are far enough away from centroids already chosen. We use a similar strategy in Kilosort2, with the added complication that spikes which are far from existing centroids would not even be detected by the online template matching step. In this case, such spikes must be detected in the residual of the model after reconstructing the data with the spikes found by template matching.

To perform these detections, we run a spike detector on the residual and pick a subset of those spikes as new templates to be introduced in the optimization. The spike detector was designed primarily to be fast, and to ensure that large amplitude spikes are not missed. It uses six single-channel prototype waveforms *w*_*k*_, *k* = 1, 2, …, 6. For each channel, we check the variance explained of all templates that extend over the nearest *n* channels with *n*≤7 and have the same single-channel waveform on each channel, chosen from one of the six single-channel prototypes *w*_*k*_. This is a much simplified version of the spike detector introduced in Kilosort2.5, but a very fast version nonetheless. Similar to the Kilosort2.5 spike detector, this detector computes the maximum variance explained at each channel and each timepoint, that can be obtained using one of the 42 template combinations described above. Using this maximum variance matrix, we find peaks that are maxima across channels for each timepoint, and then we find the subset of those which are also maxima across time, and in a neighborhood of ±4 channels in a single batch. The reason for taking the maxima across time is to ensure that no spike from the same neuron is detected twice, because these spike detections are introduced as new putative templates.

New templates are introduced on every batch. The raw data snippets at the detected spikes are first smoothed with three principal components before being added to the set of active templates. The number of spikes detected by each template is monitored using an exponential filter with a decay scale of ∼ 20 batches. Every five batches, templates are triaged and removed from the active set if their firing rate is below 0.02 spikes/s. During this step, templates are also merged together if they have a high correlation (*>*0.9) and if their means are similar (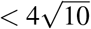 difference). If a merge is performed. the template with the smaller firing rate is simply dropped out of the active set.

#### Bimodality splits

Merges are relatively easy to perform, for example by checking all pairs of correlated templates and computing their cross-correlograms to find whether it is refractory (as described above for Kilosort4). Splits however are much more difficult, because finding a good split of a cluster in high-dimensional space is in itself a combinatorially difficult problem. In fact, the problem of finding good splits is not so different from the original problem of clustering the data, with the distinction that a split is a separation into only two, rather than many, clusters. Since we only need to divide the data into two clusters, we can take advantage of a common intuition that human operators have when performing splits: if a projection axis in the data exists which has a bimodal distribution, that is strong evidence that a split should be performed along that axis. This split can also be tested with respect to refractoriness of the CCG (like we check merges), and if the CCG is not refractory, then the split is typically performed.

How could we find such splits automatically? Human curators typically find the splits in a GUI like Phy, by investigating multiple scatter plots of pairs of principal components from a few neighboring channels. Clearly this can be improved on, since the optimal split should include information from all channels and all principal components. In Kilosort2, we designed an algorithm called bimodal pursuit to find projection axes that are highly bimodal. The “pursuit” part is a reference to other pursuit-type algorithms like ICA (kurtosis/skewness pursuit), and refers to the iterative nature of the algorithm which sequentially increases the bimodality of a projection *w*^*T*^ **x**, where *w* is the unit-norm projection vector, and **x** is the data to be split into two clusters.

Specifically, we find the projection *w* which maximizes the following log-likelihood function:

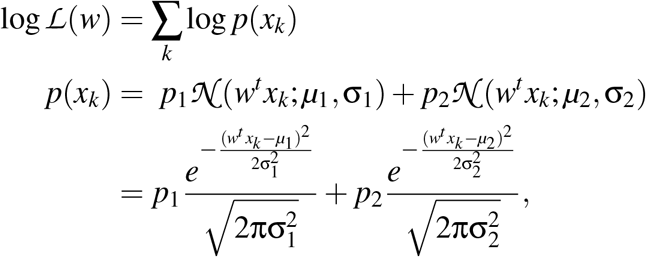

where *x*_*k*_ are the features of the *k*-th spike, *µ*_*j*_, σ_*j*_ are the scalar mean and variances of one cluster *j* out of the two total, *p*_*j*_ is the prior probability of drawing a spike from that cluster with *p*_1_ + *p*_2_ = 1. This function can be optimized via an EM-style algorithm for mixtures of Gaussians, where we first infer the posterior distribution over cluster assignments, and then we optimize the free energy function with respect to *p*_*j*_, *µ*_*j*_, σ_*j*_ given *w* and viceversa. The posterior distribution over cluster assignments is given by the “re-sponsibilities” *r*_*k j*_ = Prob(*y*_*k*_ = *j*|θ), where *y*_*k*_ is the true hidden label of data point *k* and θ is the set of all parameters.

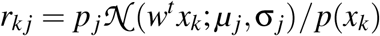

and the free energy function takes the form

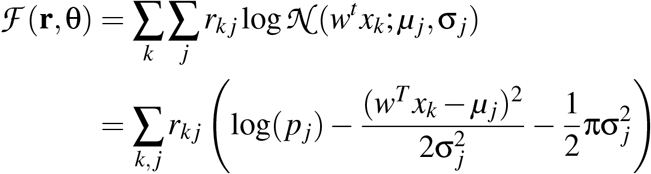

Holding *w* fixed, we can maximize with respect to *p*_*j*_, *µ*_*j*_, σ_*j*_

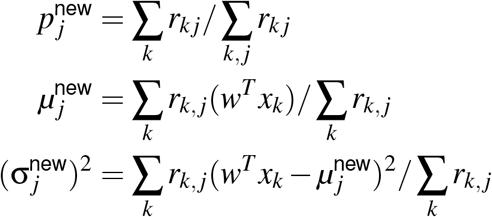

Holding *p*_*j*_, *µ*_*j*_, σ_*j*_ fixed for all *j*, we can maximize with respect to *w*:

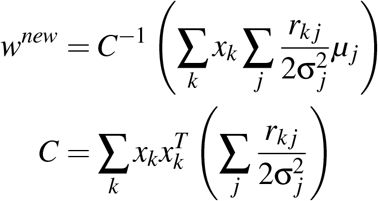

*w* is re-normalized to unit norm on every iteration. The algorithm is initialized with *w* being either the top principal component of **x**, or its normalized mean. In Kilosort 2 and 2.5, we run the algorithm twice, first initialized with the top principal component, and then initialized with the mean. The EM algorithm is run for 50 iterations, but *w* is only updated after iteration 10, and on odd iterations only, in order to make the optimization faster. We assign each spike *k* to the cluster *y*_*k*_ with highest posterior probability 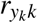. We also compute a measure of the certainty in assigning *y*_*k*_ as: 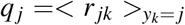. If *q*_*j*_ is very close to 1, it means all spikes in cluster *j* are assigned with nearly maximum confidence. If it is close to its lower boundary of 0.5, it means there is almost no difference between the means and variances *µ*_*j*_, σ_*j*_ of the two Gaussians in the mixture. To perform a split, we require that min(*q*_1_, *q*_2_) *>* 0.9. In addition, we require that the resulting clusters have templates that are sufficiently distinct (correlation *<*0.9 or norms *n*_1_, *n*_2_ that are sufficiently different: ∥*n*_1_ − *n*_2_∥ */*(*n*_1_ + *n*_2_) *>* 0.1). We also require that the smallest cluster in the split should have at least 300 spikes, and that the cross-correlogram between resulting clusters is not refractory, using similar criteria to those described above for Kilosort4.

Splits are performed by traversing the list of clusters in consecutive order across channels. Once a split is found, we also check the subclusters for potential splits. We do this by appending the smallest sub-cluster to the end of the list, and testing the large cluster for splits again. This process continues until no more good splits are found, and then the process moves to the next cluster in the list.

Merges are also performed at the end in all versions of Kilosort starting with Kilosort2, and they take the some form as the global merges described above in the Kilosort4 section.

### Recursive pursuit (Kilosort3)

In Kilosort3, we realized that the cost function above has some major weaknesses, such as the lack of scale invariance which means that projections with small amounts of variance have undesirably large values of log *ℒ*(*w*). Nonetheless, in practice the maximization of log *ℒ*(*w*) does indeed find projections with substantial bimodality, which is perhaps a consequence of good initialization and local minima. In Kilosort3, we made some appropriate modifications to the cost function as well as to the initialization to further improve its performance. The improved bimodal pursuit algorithm was able to find surprisingly good splits, even when given a mixture of more than two clusters. We took advantage of its performance and designed a new clustering algorithm in Kilosort3 which performs clustering by recursively splitting off clusters from the main distribution using the bimodal pursuit algorithm.

#### Improved bimodal pursuit

The idea of “projection pursuit” comes from the field of independent components analysis (ICA) and similar algorithms, and it was perhaps popularized the most by the fast ICA algorithm from Hyvarinnen and colleagues [39]. Like our cost function, projection pursuit maximizes some criterion computed over the distribution of projections *w*^*T*^ *x*. Unlike our function, this criterion is usually scale-invariant, such as the kurtosis or skewness criteria which are normalized by the variance of the data. A simple way to make our criterion scale-invariant is whitening or “sphering”, which is also a very common preprocessing step for ICA. Whitening normalizes a multi-dimensional dataset **x** into 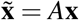, where *A* is an appropriate whitening matrix, so that the mean of each dimension is 0, and the covariance of the normalized data is the identity. As a consequence, any projection 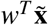 is standardized, in other words it has mean 0 and variance 1. A common choice of whitening is PCA / SVD, and this is also our approach.

In Kilosort3, we perform whitening on every matrix **x** before running the bimodal pursuit algorithm. In this case, the log-likelihood criterion can be interpreted as searching for the projection *w* which can be best modelled by a mixture of Gaussians after z-scoring. Of all distributions with mean 0 and variance 1, the criterion *L*(*w*) is now maximized by the sum of discrete distributions centered on − 1 and +1. In addition, we constrain σ_1_ = σ_2_ = σ, which allows to perform the matrix inversion *C*^−1^ only once, because

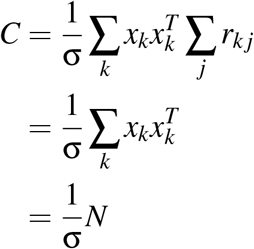

because ∑ _*j*_ *r*_*k j*_ = 1 by construction and 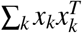 due to whitening, where *N* is the total number of spikes.

We also introduce a new form of initialization. Since the algorithm is highly sensitive to initialization, we run a brute force search for a good initialization vector *w*. Remembering that we are in normalized PCA space, the brute force approach checks all vectors *w* with *w*_*d*_ ∈ {−1*/n*, 1*/n*} for *d* = 1, …, 6 and *w*_*d*_ = 0, *d >*6, with *n* being a normalization constant, in this case 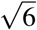. For each of the resulting 64 combinations, we check the bimodality of the projection *w*^*T*^ **x** by performing a histogram and using similar criteria to those described above for Kilosort4. This histogram based algorithm is also used in the first 25 iterations of bimodal pursuit as a replacement for the EM assignments for µ, σ, *p*. This is done by computing the mean, variance and fraction of all points *w*^*T*^ *x*_*k*_ smaller/bigger than the trough of the distribution respectively. We found this initial approximation of the EM assignments to be more robust, especially in cases where one cluster has substantially more spikes than the other.

#### Recursive pursuit

The bimodal pursuit algorithm described above takes as input a set of spike features, and outputs a partition into two clusters. Applied recursively, the algorithm can find a subset of a dataset that is well isolated from other clusters and cannot be split further into more clusters. We start will all spikes detected on a set of channels, and find the first split. Of the two pieces, we take the piece with higher average waveform amplitude, and we split it again. This process continues until no more splits can be found. Splits can be veto-ed in similar ways to Kilosort 2/2.5, except that the index of bimodality is used instead of the “measure of certainty” described above. A split requires all three criteria to be satisfied: high bimodality index, low waveform correlation and non-refractory CCG. In Kilosort4, we use the same criteria minus the criterion for low waveform correlation which is somewhat redundant with the bimodality index criterion.

Once a cluster is found out of the dataset, the spikes corresponding to that cluster are removed, and the cluster finding process is applied to the remaining spikes. This process continued until the remaining spikes can no longer be split, and thus they constitute the final cluster. Note that the complete algorithm contains two recursive loops: one loop for finding a single cluster out of the dataset, and another loop for finding all clusters in the dataset. This clustering operation is applied to spikes detected in 40 µm segments of the probe, similar to the process described for Kilosort4.

### Benchmarking

To determine the performance of various spike-sorting algorithms, we created realistic simulations using the properties of 512 electrophysiological recordings from the International Brain Laboratory (IBL) performed using Neuropixels 1.0 probes [15, 40]. These recordings were processed by the IBL using pyKilosort. The simulation generation was over two times faster than realtime (e.g. a 45 minute simulation took around 20 minutes to generate), which enabled us to create several simulations for benchmarking. The simulations, other than “step drift, aligned”, used the site configuration of the Neuropixels 1.0 probes, which have a vertical spacing of 20 µm, a horizontal spacing of 32 µm and a horizontal offset across rows of 16 µm.

#### Drift simulation

pyKilosort, like other Kilosort versions, returns the estimated depth for each processing batch at nine equally-spaced positions along the 3.84 mm probe. The processing batch size for all IBL recordings was 65,536 timepoints. We quantified the drift range for each recording by first taking the median of the depth across the nine positions, then computing difference between the 5^th^ and 95^th^ percentile of the drift. We used the properties of the drift across these recordings to create simulated drift (see drift examples in Figure S3).

For the simulations, we generated a drift trace of length 45 minutes at each of the positions, then up-sampled the drift to all 384 channels using linear interpolation. The drift was the same across a period of 2 seconds for all simulations, other than the fast drift simulation which varied in periods of 200 ms. Here are the details of the generation of each drift simulation:

- *no drift* : Zero drift at all nine positions.
- *medium drift* : The overall drift was generated as random gaussian noise smoothed in time with a gaussian filter of σ=100 sec. Drift at each of the nine positions was generated as random gaussian noise smoothed in time with a gaussian filter of standard deviation 100 seconds and smoothed across the positions with a gaussian filter with σ=2. This perposition drift was rescaled by a factor of 0.4 and added to the overall drift, then the drift across positions and time was rescaled such that the minimum and maximum values were -7 µm and 7 µm. This resulted in a simulation with a drift range of 9.4 µ.
- *high drift* : The overall drift and per-position drift were generated in the same way as the medium drift. The per-position drift was next rescaled by a factor of 0.26 and added to the overall drift, then the drift across positions and time was rescaled such that the minimum and maximum values were -18.5 µm and 18.5 µm. This resulted in a simulation with a drift range of 27.9 µm.
- *fast drift* : A medium drift simulation was used for the slow drift across positions and time (generated in bins of two seconds, then upsampled to 200 ms bins with nearest neighbor interpolation). Then fast drift events were generated with an amplitude of 10 µm and a difference of exponentials kernel with a rise time of 80 ms and a decay time of 200 ms. 300 of these fast drift events were added to the upsampled medium drift simulation at random times.
- *step drift* : The overall drift and per-position drift were generated in the same way as the medium drift. The per-position drift was next rescaled by a factor of 0.58 and added to the overall drift, then the drift across positions and time was rescaled such that the minimum and maximum values were -4 µm and 4 µm. Halfway through the recording 30 µm was added to all the drifts across positions.
- *step drift, aligned* : Same exact drift as step drift, but waveforms were upsampled using aligned probe sites with a vertical separation of 20µm and a horizontal separation of 32µm.

#### Extraction of waveforms at multiple depths

Obtaining waveforms across many depths requires recordings with substantial drift. In the IBL dataset we found 11 such recordings with high drift that well-sampled a range of 40 µm in depth. We used the estimated spike times from pyKilosort for each detected unit in these recordings, and the estimated depth of the probe to compute the average waveform for the unit at specified depth positions. We used 20 depth bins each of size 2µm, resulting in average waveforms across 40 µm. To ensure the quality of the waveforms, we did not use any units which had fewer than 50 spikes at each depth.

The waveforms were denoised by reconstructing each waveform across depths with only its top three principal components. The waveforms were then normalized by the average norm of the waveform across depths. We then threw out waveforms which varied substantially from -20 µm to 20 µm in depth, since these waveforms’ shape changes are likely caused by other processes besides drift. To quantify the variation across depth we computed the Euclidean distance across channels and timepoints between the waveform at -20µm and the waveform at 20µm shifted up by 4 channels (a distance of 40 µm). We removed units with variation greater than 0.25 (∼ 25% of units), resulting in a waveform bank of 597 units from the 11 recordings.

Next we needed the waveform shapes at a finer scale than 2µm. For this, we upsampled the waveforms by a factor of 100 using kriging interpolation [9] with a regularization coefficient of 0.01 and a gaussian of standard deviation 20µm. For the step drift with aligned site simulation, the upsampled waveforms were interpolated using a probe with sites aligned vertically. Then the waveforms were again normalized by the average norm of the waveform across depths. We next divided these waveforms into two groups according to the contamination rates from their units’ estimated spike trains [41]: a contamination rate less than were used to generate “single-units”, while those with a contamination rate greater than 0.1 were used to generate “multi-units”.

The units from these recordings exhibited waveform changes across depth, see example waveforms in Figure 3c and Figure S2a. All waveforms moved down the probe as the depth changes, but some waveforms also changed their shape (example units 2 and 4, which had smaller spatial footprints). This shape change could not be inferred by other channels. We demonstrated this by using the same Kriging interpolation procedure as above in order to estimate the 0 µm depth waveform from the waveforms at other depths (Figure S2b). The waveform at 0 µm depth was well-estimated for units 1 and 3 but not for 2 and 4. This exemplifies the need for real recordings to create accurate simulations of waveform shapes.

We quantified the performance of the spike-sorting algorithms as a function of the spatial extent of the waveforms of the ground truth unit (Figure S5c,f). We defined the spatial extent of the waveform as the spatial scale across channels over which the waveform shape is maintained (using the 0 µm depth waveform). To compute this we first matched the waveform with its most similar waveform from the universal templates as defined by cosine similarity. We then projected the waveform onto this best template waveform and thresholded it, to obtain a template weight for each channel. We next computed the weighted mean of the distance from each channel to the center-of-mass of the waveform as defined by the template weights, and termed this the spatial extent.

#### Simulation of spikes

We simulated 600 “single-units” and 600 “multi-units” by randomly drawing waveforms from these two classes. These waveforms were randomly placed on the probe at positions from site 4 to site 380. To create the correct waveform shapes, the waveform’s best channel modulo 4 was computed and maintained in the simulation (because the probe site arrangement repeats every 4 sites).

We used the inter-spike intervals from detected units in the 11 recordings which had a contamination rate of less than 0.1, this was 1497 units in total. The average firing rate of these units was 12.6 Hz. Each simulated spike train for a “single-unit” was then generated by randomly shuffling the inter-spike intervals of one of the detected units. For the spike trains of “multi-units” we generated Poisson spike trains with firing rates drawn randomly from these units’ firing rates.

The amplitudes for the “single-units” were generated by adding a constant, 10, to a random exponential with a mean of 7, which approximated the distribution from units detected in the data. The amplitudes for the “multi-units” were generated from a uniform random distribution with a range from 4 to 10. The waveform across depth for each unit was then multiplied by its amplitude. We quantified the performance of the spike-sorting algorithms as a function of the amplitude (norm) of the ground truth unit (Figure S5b,e).

We then added one-by-one the spike train of each simulated unit to the simulation, using the simulated drift at each timepoint to determine which depth of the waveform to add for each spike. Collisions could occur in the spike trains, so we added the spike train in 3 interleaved parts to ensure correct reconstruction while still maintaining the speed of simulation generation.

All simulations used different waveforms, spike trains, and amplitudes, except for the two step drift simulations, in which all parameters were kept fixed to determine the effect of probe site configuration. These two step drift simulations therefore only differed in their exact waveform shapes across depths due to the difference in the probe site positions.

#### Simulation noise and “unwhitening”

To each channel in the simulation we added random noise with a flat frequency spectrum in time, up to 300 Hz. This noise was scaled to have a standard deviation of 0.76. Next the simulation was “unwhitened”: the simulation was multiplied by the inverse of a whitening matrix estimated from one of the 11 recordings used. Different whitening matrices were used for each simulation, except for the two step drift simulations, where it was the same matrix for both. Finally, to save the simulation as int16, the simulation was multiplied by 200, cut-off at ± 32767 and converted to int16. For each simulation we saved a corresponding “.meta” file, which SpikeInterface expects for processing IMEC Neuopixels probe recordings. For the aligned site probe, we added a probe type to the spike GLX loader in SpikeInterface.

#### Performance metrics

Each ground truth unit was compared to the 20 closest detected units from the algorithm, where closeness was defined by the distance between the ground truth and detected units best channels. If an estimated spike from a detected unit was less than or equal to 0.1 ms from a ground truth spike, it was counted as a positive match. The false positive rate (FP) was defined as the number of estimated spikes without a positive match divided by the total number of estimated spikes, which is equivalent to 1 − *precision*. The false negative rate (FN) was defined as the number of missed ground truth spikes divided by the total number of ground truth spikes, which is equivalent to 1 − *recall*. We matched the ground truth unit with the detected unit that maximized the *score*, defined as 1 − *FP* − *FN* [6]. The upper bound of the score is 1. In Figure 3e-j, the ground truth units were sorted by their score from each algorithm separately. We defined ground truth units as being correctly identified in Table 2 if the score was higher than 0.8.

#### Spike-sorting algorithm parameters

We ran Kilosort 1, 2, 2.5 & 3, IronClust, Mountain-Sort4, SpyKING CIRCUS, SpyKING CIRCUS 2, HD-Sort, Herding Spikes, and Tridesclous2 on all simulations using the SpikeInterface platform to ensure that all spike-sorting algorithms were run in the same way. For Kilosort 1, 2, 2.5 and 3, we set the detection thresholds to [9, 8] instead of their defaults which varied across verrsions. Also, to speed up Kilosort 1, we set the number of passes through the data to 2 instead of 6 (this did not reduce performance).

For the other top performing algorithms (IronClust, MountainSort4, and SpyKING CIRCUS), we ran a parameter sweep over the detection threshold and used the detection threshold which maximized the number of correctly identified units on the medium drift simulation. For MountainSort4 and IronClust, the best detection threshold was the default detection threshold; for SpyKING CIRCUS, this was a detection threshold of 4.5. For SpyKING CIRCUS 2, we noticed poor detection of low amplitude units (Figure S5b,e) and thus also swept over the detection threshold for this algorithm, but did not achieve an improvement in performance. For IronClust the default adjacency radius is 50, while for MountainSort4 the default is set to all channels. This large radius led to an incredibly long run time (10s of hours), and thus we set the Mountain-Sort4 adjacency radius to 50 as well.

All other parameters were set to their default values.

#### Comparison to other benchmarking approaches

Here we compare our approach to previous spike-sorting benchmarking performed in the literature. The first approach is to use datasets where the ground-truth spiking of a single unit is known. These datasets are acquired by performing cell-attached recordings while simultaneously recording with a probe. Then spike-sorting is performed on the probe and compared to the ground-truth spiking to determine spike-sorting performance. Since these are very difficult experiments, existing ground-truth datasets were acquired in anesthetized animals and are very short [2, 42–48]. This makes these datasets much easier to spike sort compared to long, realistic awake recordings with drift and with relatively more neuronal firing. Further, such cell-attached recordings are biased towards larger and higher firing units, and less likely to be successful on smaller amplitude units. When SpikeForest used these ground-truth datasets to compare various spike-sorting algorithms (“PAIRED” recordings, https://spikeforest.flatironinstitute.org/) [49], they found that IronClust, Kilosort2 and SpyKING CIRCUS performed similarly on these recordings. This is consistent with our own benchmarking results on the “no drift” recordings, where many of the spike-sorting algorithms recovered units with high amplitudes equally well (Figure S5b,e). However, most recordings in awake animals have drift and contain many low amplitude units that can be isolated by Kilosort.

Another approach is to create so-called “hybrid ground-truth datasets”. Either ground-truth units, as acquired above, or manually curated units are used [6, 8, 49]. These units can be inserted into other real recordings, or the same recording in a different position after being subtracted off, to ensure appropriate background noise. Multiple ground truth units can be inserted in a dataset in this fashion. However, these hybrid datasets depend on finding the neurons in the first place and they also depend on correcting for the initial drift of the dataset. Alternatively, these ground-truth units can be used to create simulations with drift, in which the units are moved up and down the probe at some rate over time, then background noise is added [46, 50]. These simulations do have drift but they have drawbacks: 1) waveform changes across depths cannot occur, but they do in the real data, as demonstrated in Figure S2a,b, and 2) the background noise is only simple gaussian noise with the same frequency spectrum as the real data. To remedy (1), we obtained waveforms at various drift positions from real recordings, as outlined above, to simulate the waveforms at various depth positions. To remedy (2), we added 600 “multi-units” with low amplitudes to the simulation to create more realistic background, on top of adding gaussian noise with a matched frequency spectrum (Figure S2c).

The final approach is to instead simulate waveforms, either using some specified properties [3], or using the electrical field of a biophysically simulated neuron [51–54]. These simulators do not produce waveforms as diverse as real neurons from recordings, likely because we lack a full understanding of how the tissue geometry interacts with action potentials and the probe to create all the diverse spike shapes that can be observed. Various types of noise and background can be added to these neurons. For example these simulated neurons can be added to background signal from other recordings [3]. Alternatively noise can be added by simulating neurons further away from the probe [53]. Other simulators use spatially correlated noise with parameters extracted from the data [51, 54]. The MEAREC simulator includes the option for probe drift; however it is unclear how much the waveform shape changes over drift positions in their simulations, as this depends on the geometry of the electrical fields.

---

**S1:**
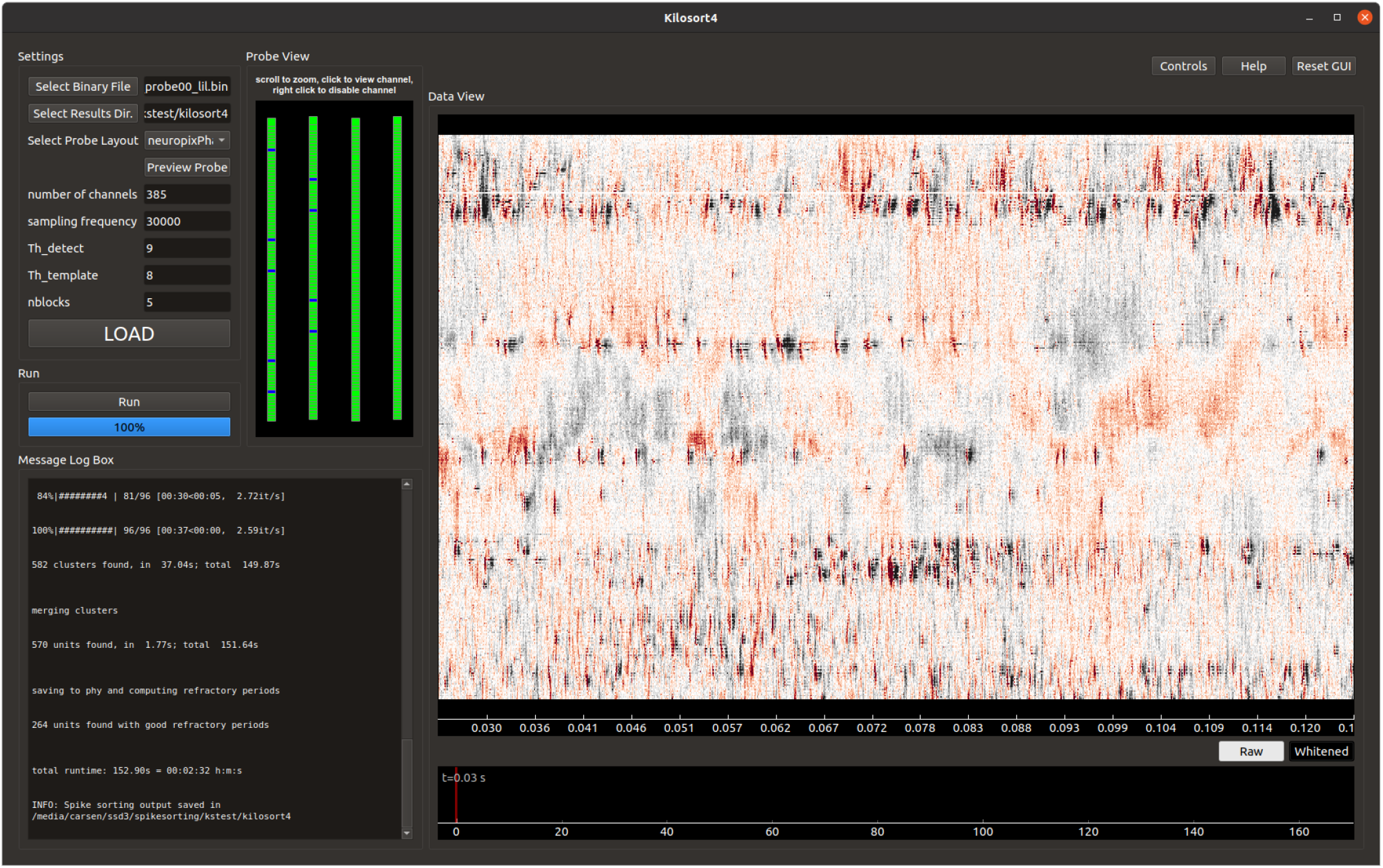
Kilosort4 graphical user interface. The GUI for Kilosort4 enables the user to load in and view the binary file both raw and whitened. Next the user runs the spike-sorting pipeline. The message log box allows the user to monitor the progress of the spike-sorting algorithm.

**S2:**
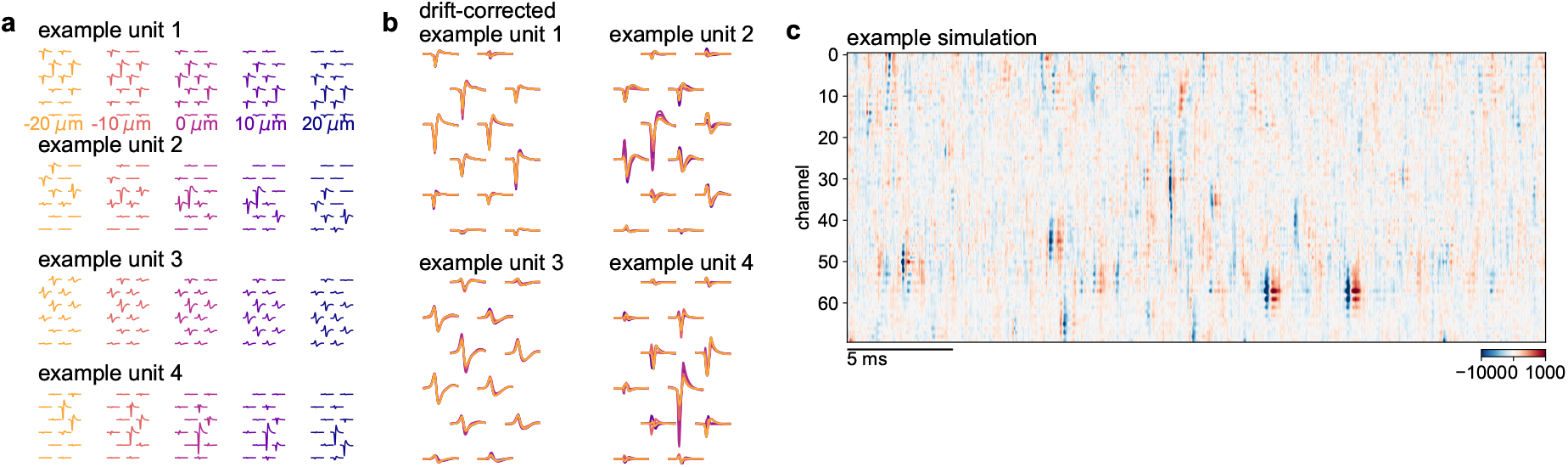
Simulation features. **a**, Four additional example units like in Figure 3c. **b**, The units in **a** after drift correction with the interpolation method from Kilosort 2.5/3/4. Units 1 and 3 interpolate well, while units 2 and 4 do not, due to having features with small spatial footprints. **c**, Example section of the simulation in Figure 3d, but after independent noise was added and the data was un-whitened.

**S3:**
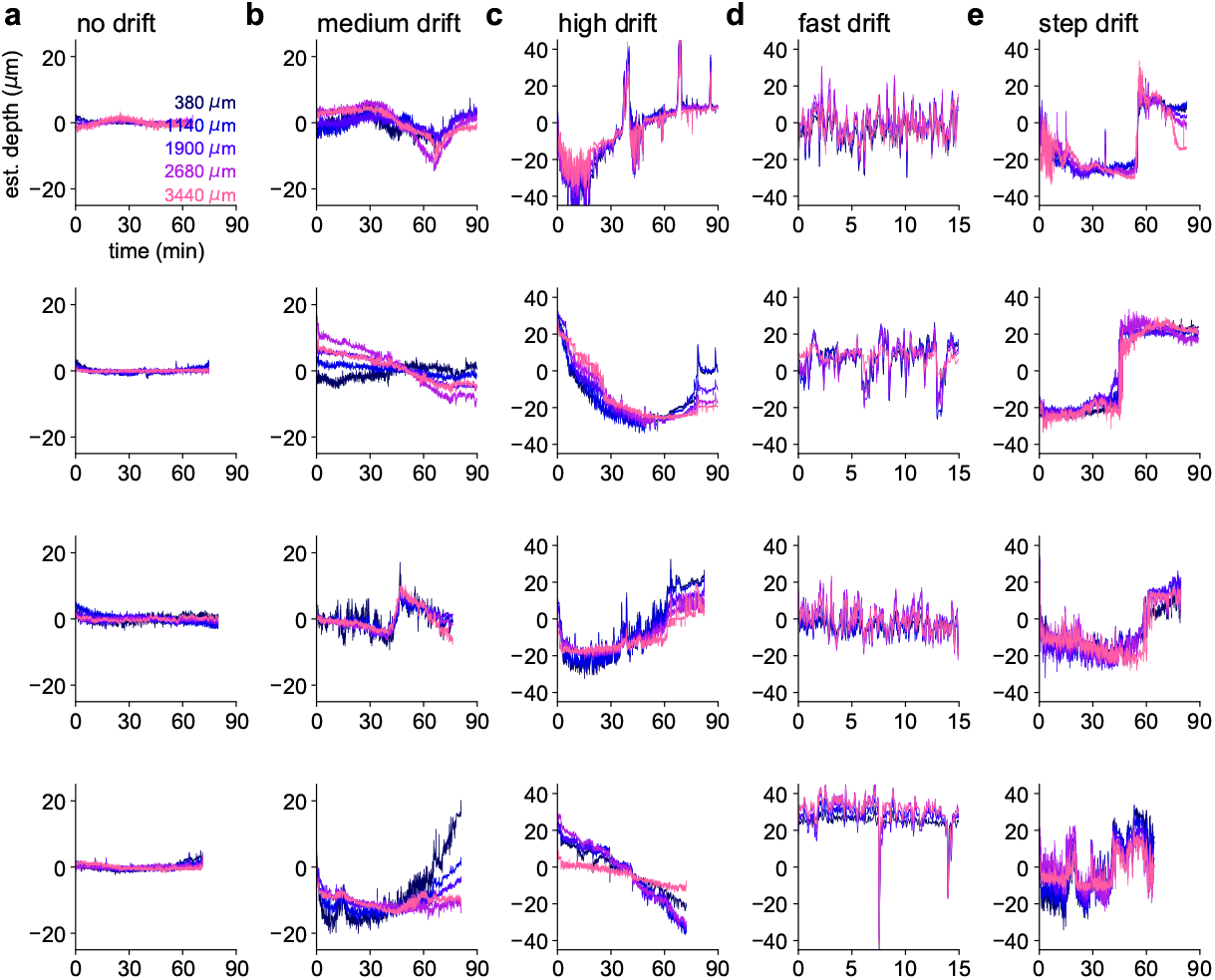
Real drift examples. These are inferred drift traces from the IBL dataset grouped into: **a**, no/small drift **b**, medium drift, **c**, high drift, **d**, fast drift and **e**, step drift. Note that in many cases different types of drift are combined.

**S4:**
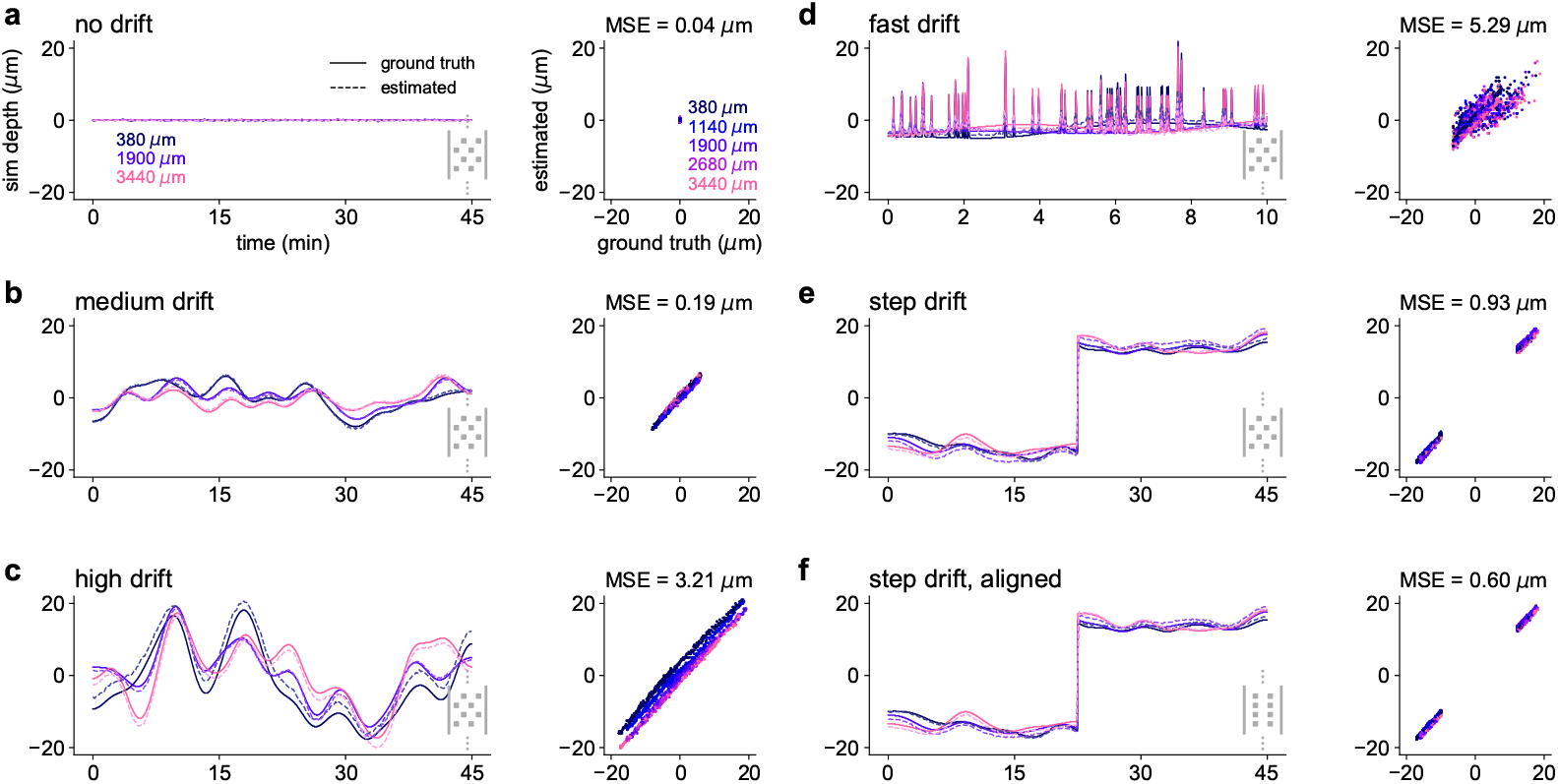
Recovered drift traces from simulations. **a-f**, Ground truth simulated drift + the drift identified by Kilosort4. (Left) Estimated and true drift traces. (Right) Scatter plot of estimated and true drift traces.

**S5:**
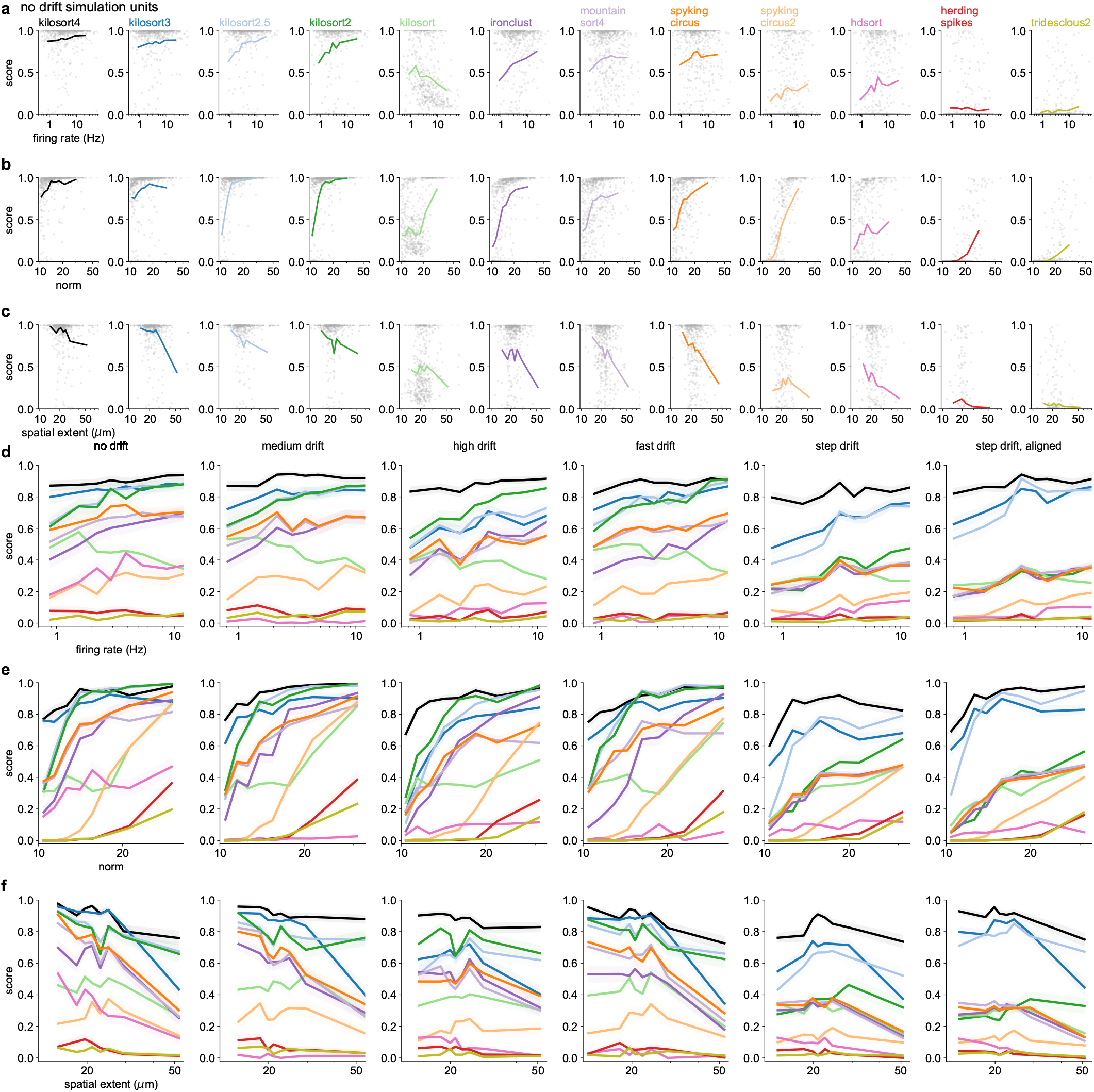
Accuracy as a function of firing rate, amplitude/norm and spatial extent. **a-c**, Scatter plots of unit properties (firing rate, norm, spatial extent respectively) versus accuracy, for the no drift simulation. Lines show the average accuracy in bins of wqual numbers of points. **d-f**, Average accuracy curves for all types of simulations and all unit properties (firing rate, norm, spatial extent respectively).

